# Bats are key hosts in the radiation of mammal-associated *Bartonella* bacteria

**DOI:** 10.1101/2020.04.03.024521

**Authors:** Clifton D. McKee, Ying Bai, Colleen T. Webb, Michael Y. Kosoy

## Abstract

Bats are notorious reservoirs of several zoonotic diseases and may be uniquely tolerant of infection among mammals. Broad sampling has revealed the importance of bats in the diversification and spread of viruses and eukaryotes to other animal hosts. Vector-borne bacteria of the genus *Bartonella* are prevalent and diverse in mammals globally and recent surveys have revealed numerous *Bartonella* lineages in bats. We assembled a sequence database of *Bartonella* strains, consisting of nine genetic loci from 209 previously characterized lineages and 121 new cultured strains from bats, and used these data to perform the most comprehensive phylogenetic analysis of *Bartonella* to date. This analysis included estimation of divergence dates using a molecular clock and ancestral reconstruction of host associations and geography. We estimate that *Bartonella* began infecting mammals 62 million years ago near the Cretaceous-Paleogene boundary. Additionally, the radiation of particular *Bartonella* clades correlate strongly to the timing of diversification and biogeography of mammalian hosts. Bats were inferred to be the ancestral hosts of all mammal-associated *Bartonella* and appear to be responsible for the early geographic expansion of the genus. We conclude that bats have had a deep influence on the evolutionary radiation of *Bartonella* bacteria and their spread to other mammalian orders. These results support a ‘bat seeding’ hypothesis that could explain similar evolutionary patterns in other mammalian parasite taxa. Application of such phylogenetic tools as we have used to other taxa may reveal the general importance of bats in the ancient diversification of mammalian parasites.

**Significance statement:** Discovering the evolutionary history of infectious agents in animals is important for understanding the process of host adaptation and the origins of human diseases. To clarify the evolution of the *Bartonella* genus, which contains important human pathogens, we performed phylogenetic analysis on a broad diversity of *Bartonella* strains, including novel strains from bats. Our results indicate that *Bartonella* clades diversified along with their mammal hosts over millions of years. Bats appear to be especially important in the early radiation and geographic dispersal of *Bartonella* lineages. These patterns are consistent with research indicating a chiropteran origin of important human viruses and eukaryotic parasites, suggesting that bats may play a unique role as historical sources of infections to other hosts.

## Introduction

A central part of the work done by disease ecologists is to understand the host range of infectious agents. However, host ranges must be understood in a coevolutionary context, specifically how agents have adapted to and diversified in hosts over time. Only by considering both ecological and evolutionary context can we understand how agents come to infect and adapt to new hosts. While cophylogeny is a common tool for studying the codiversification of hosts and parasites, few studies have examined the relative timing of the diversification of parasite lineages in parallel with that of hosts (1, 2).

The genus *Bartonella* is an excellent study system for disease ecology and evolution because it is common and diverse in many mammalian hosts (3). These alphaproteobacteria are facultative intracellular pathogens that can cause persistent, hemotropic infections in their hosts. Transmission between hosts occurs through a variety of hematophagous arthropod vectors, wherein bartonellae colonize the midgut and are then shed in arthropod feces (4). Clades of *Bartonella* species tend to be host-specific (5), so it could be hypothesized that the genus diversified along with its mammalian hosts millions of years ago. However, there have been few comprehensive phylogenies of this genus and limited research on the influence of particular host groups on *Bartonella* evolution.

Bats are a group of special interest because they have traits that are amenable to parasite transmission, including their global distribution, ability to fly, seasonal migration, dense aggregations and high sociality in some species, long life spans, and the use of torpor and hibernation (6). There is also evidence that chiropteran immune systems are highly tolerant of infections, especially of viruses (7). Thus, their role as reservoirs for *Bartonella* bacteria may be uniquely influential among mammals. Bats are also an ancient clade of mammals (8), providing ample time for diversification of bacterial parasites and transitions from bats to other mammals. Research has concluded that bats are potentially ancestral hosts that influenced the diversification and spread of coronaviruses (9), lyssaviruses (10), paramyxoviruses (11), trypanosomes (12), and haemosporidia (13) among other mammalian orders. Drawing from Hamilton *et al*. (14), who developed the ‘bat-seeding’ hypothesis to explain the geographic and host distribution of *Trypanosoma* lineages related to the agent of Chagas disease, *T. cruzi*, we hypothesize that bats may have also been influential in the ancient diversification and spread of *Bartonella*.

Successful amplification of *Bartonella* DNA from recent fossils points to a prolonged history of *Bartonella* infection in some hosts, such as humans and domestic cats (15). However, it is unlikely that DNA could be successfully amplified from more ancient fossils to test hypotheses about the origin of bartonellae in mammals. Instead, a molecular clock approach can be used to estimate the rate at which substitutions accumulate in *Bartonella* DNA and then extrapolate divergence dates of clades. Considering new research has shown that mammal-associated bartonellae evolved from arthropod symbionts (16), we rely on a molecular clock for the 16S ribosomal RNA (rRNA) gene based on sequence divergence data from bacterial symbionts of arthropod hosts separated for millions of years (17). We perform a multi-locus analysis using the most comprehensive database of *Bartonella* strains to date, including a greater number of loci than a recent time tree analysis (18) and broader taxon sampling than previous genomic analyses (19). Many new *Bartonella* strains have recently been discovered in bats (20), so we have included 121 novel strains of bats in this study to amend current delineation of *Bartonella* clades (19) and to determine the influence of bats on the diversification and spread of *Bartonella* bacteria to other mammalian orders.

Using this molecular clock approach, we extrapolate when the genus *Bartonella* diversified and compare the timing of *Bartonella* clade diversification along with their hosts. We hypothesize that mammal-infecting bartonellae evolved with their hosts starting in the late Cretaceous or early Paleogene when many eutherian and metatherian taxa diversified (21). We expect to see clustering of lineages associated by host orders and correlation between diversification dates of hosts and *Bartonella* clades. Using ancestral state reconstruction and network analysis, we discern which orders of mammals were highly influential in the diversification and spread of *Bartonella* to other host orders and geographic regions. We predict that the speciose orders of bats (Chiroptera) and rodents (Rodentia) are important in the historical expansion of the *Bartonella* genus, however bats may have a more profound influence in this process because of their ability to fly and quickly disperse over wide areas. This study provides a more complete understanding of *Bartonella* evolution and biogeography and the role of bats as important hosts of pathogens through a suite of phylogenetic methods that can be adapted to understand these processes in other host-specific parasites and symbionts. Such investigations could lead to a deeper evolutionary understanding of symbiosis and parasitism and the identification of key host groups in the diversification and spread of these organisms.

## Materials and methods

### Molecular data collection

To balance the need for increased taxon sampling and adequate sequence data to produce a well-supported phylogeny, we assembled a database of *Bartonella* sequences from published genomes on GenBank, previous studies using multi-locus sequence analysis (MLSA), and archived cultures from bats. We targeted nine genetic markers (SI Appendix, Table S1) commonly used for *Bartonella* detection and phylogenetic analysis (22). Data from MLSA studies and genomes published as of 2018 were collected from GenBank via accession numbers or strain numbers from 74 studies (SI Dataset 1), including recent publications that have isolated bartonellae or related bacterial symbionts in arthropods and past studies characterizing bat-associated *Bartonella* strains from Asia, Africa, and North America. We excluded any strains that were noted in the studies as showing evidence of homologous recombination between *Bartonella* species to prevent issues with incomplete lineage sorting in phylogenetic analysis. Additional molecular data collection of *Bartonella* strains from bats included a subset of cultures archived in our laboratory from previous studies in Africa, North and South America, Europe, and Asia that have been partially characterized at some of the targeted loci, as well as new cultures from bats sampled from Nigeria in 2010 and Guatemala in 2010, 2014, and 2015. The data combined from bat-associated *Bartonella* strains cover 50 species from 10/20 extant chiropteran families (8). Details on sequencing of bat-associated *Bartonella* strains, alignment, data cleaning, and validation can be found in SI Appendix. The final database contained sequence data from 332 taxa: 209 *Bartonella* reference strains from genomes and MLSA studies, 121 bat-associated strains from our laboratory archive, the ant symbiont *Candidatus* Tokpelaia hoelldoblerii, and the outgroup *Brucella abortus* (SI Datasets 1 and 2).

### Phylogenetic analysis

Bayesian phylogenetic analysis was performed using BEAST v1.8.4 (23) via the CyberInfrastructure for Phylogenetic RESearch (CIPRES) Science Gateway portal v3.3 (24). The nine loci were analyzed separately using GTR+I+G sequence evolution models, estimated base frequencies, four gamma rate categories, an uncorrelated relaxed clock with an exponential distribution of clock rates along branches for each locus, and a birth-death speciation model with incomplete sampling (25). *Brucella abortus* was set as the outgroup in all analyses. To determine a clock prior for the 16S rRNA locus, we analyzed published 16S rRNA sequence divergence and host divergence times for bacterial symbionts of arthropods (17). A linear regression model was fit to the data in R (26) and a lognormal prior was estimated by moment matching to the normal distribution for the fitted mean and standard error of the slope (SI Appendix, Fig. S5). The prior distribution for the exponential clock rate for 16S rRNA was set to this lognormal distribution while prior distributions for the exponential clocks of the remaining eight loci were set to an approximate reference prior for continuous-time Markov chain (CTMC) rates (27). Thus, the 16S rRNA clock acts a strong prior and the rates for the other eight loci are estimated relative to the 16S rRNA rate. This approach allows for external validation of *Bartonella* diversification events based on host diversification dates without explicitly using host diversification dates as calibration points for the parasite tree. Extensive testing using alternative substitution (with or without codon partitioning), clock, and tree models and subsets of genetic data determined that model choice or the exclusion of the ITS locus had little influence on tree topology and estimated divergence dates (SI Appendix, Table S5). Additional details regarding model priors and run settings can be found in SI Appendix.

### Ancestral state reconstruction

In addition to divergence time estimation, we performed ancestral state reconstruction in BEAST. We assigned discrete traits to each tip based on the taxonomic order of the host and the ecozone (28) that includes the majority of the host’s geographic range. The association of some *Bartonella* lineages with arthropods and not mammals are justified in SI Appendix. Ancestral state reconstruction was performed using a symmetrical rate model to reduce the number of state transitions that needed to be inferred.

### Tip-association tests

We performed tip-association tests using the Bayesian Tip-association Significance testing (BaTS) program v1 to assess the clustering of traits along tips of the phylogenetic tree (29). We performed four sets of simulations using the same assignments of host orders and geographic ecozones used in the ancestral state reconstruction above. The two sets of traits were simulated on 1000 posterior sampled trees from the final BEAST run and on the single maximum likelihood (ML) tree. Clustering of traits was measured by the association index (AI) and parsimony score (PS), producing a distribution for the 1000 Bayesian trees and a single value for the ML tree. Null distributions for these measures were generated using 100 randomizations of traits onto tips of the trees. The significance of clustering was evaluated based on the overlap between observed values or distributions of AI and PS and their null distributions. For both measures, small values indicate a stronger phylogeny-trait association (29).

### Host clade definitions and divergence dates

We defined host-associated *Bartonella* clades *a posteriori* based on high posterior support (>0.9) and clustering by host orders from the ancestral state reconstruction (Fig. 1A). Previous analyses of *Bartonella* host associations have shown that host-switching is common (30), so a calibration approach that assumes strict cospeciation across the tree would not accurately reflect the evolutionary history of these bacteria. However, *Bartonella* lineages are broadly host-specific within orders (18) and host-switching is more frequent between closely related hosts (31). We defined 15 host-associated *Bartonella* clades (Tables S5-S6) at relevant taxonomic scales below the order level to test the hypothesis that *Bartonella* lineages diversified with their hosts while accounting for frequent host-switching that could occur within a host clade. We collated divergence dates for the most recent common ancestor uniting the host taxa of interest within each clade from available studies in the TimeTree database (http://timetree.org/), summarized by the estimated mean, 95% confidence intervals, and range of dates across studies (32). We then correlated these mean host divergence dates with our estimated median divergence date of the associated *Bartonella* clade (Table S7). A significant linear fit between these dates would support the hypothesis that *Bartonella* diversified within their hosts after colonization. To validate measurement of the divergence time for mammal-associated *Bartonella* clades with the ultrametric tree produced in BEAST, we also generated a calibrated timed phylogeny with the ML tree. Using the RelTime relative rate framework (33) within MEGA v10.0.5 (34) we generated a timed phylogeny using host clade divergence dates from TimeTree (Table S7). We used confidence intervals (or ranges in the case of clade J) for the 15 host clade divergence dates as minimum and maximum divergence dates in RelTime. The program then calculated divergence dates on the tree using a maximum likelihood approach (33), producing mean estimates and 95% confidence intervals for clade dates that we could compare with the eubartonellae date estimated in BEAST. This analysis can confirm that the date estimation is robust to different approaches by comparing a calibration-based method on an existing tree to a method that relies on relaxed clock priors during tree estimation.

**Fig. 1.**
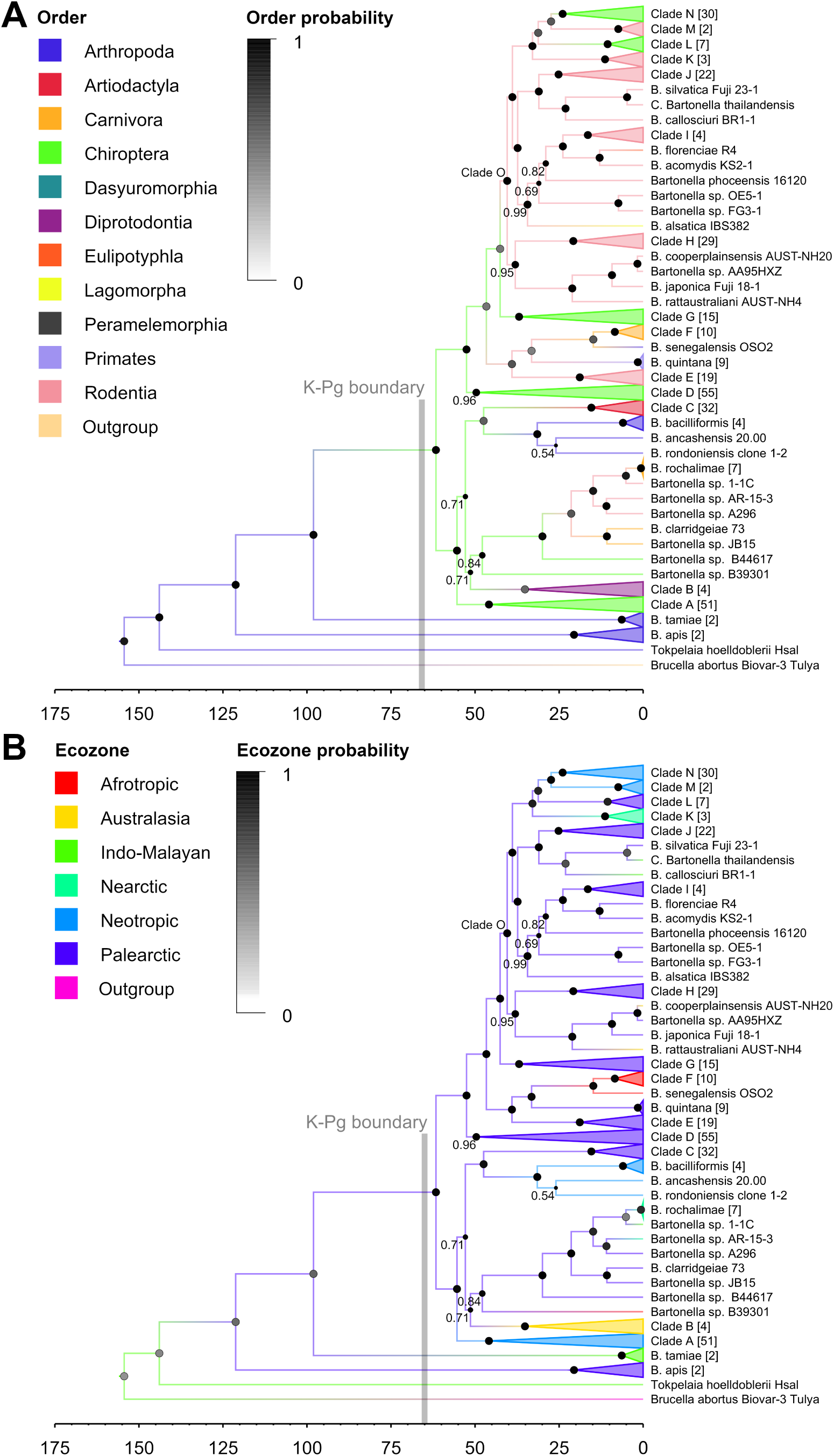
Evolution of *Bartonella* lineages, host associations, and geographic origins. Timed maximum clade credibility tree of *Bartonella* lineages. Tips are collapsed into clades of related *Bartonella* species and strains (SI Appendix, Tables S5 and S6); the number of tips in each clade is shown in brackets. Posterior probabilities (PP) for nodes are indicated by the size of circles; ancient nodes had strong support (PP = 1), unless otherwise labeled. Branch lengths are in millions of years. Ancestral state reconstruction of (A) host order and (B) ecozone transitions was performed during the estimation of diversification times. Branches are colored according to their most probable (PP > 0.5) host order or ecozone states, with host or ecozone probability shown by the color of circles at each node. The Cretaceous-Paleogene extinction event is drawn as a gray line at 66 million years ago.

### Stochastic character mapping and network analysis

To determine the inferred ancestral host order and ecozone of mammal-infecting eubartonellae, we initially inspected the results of the ancestral state reconstruction on the maximum clade credibility (MCC) tree. Specifically, we inspected the posterior support for the node and the posterior probability of the host order and ecozone at the node across all posterior trees. However, due to the large number of *Bartonella* lineages associated with Chiroptera in the database (n = 160) relative to those in other diverse orders (Rodentia, 87; Artiodactyla, 32; Carnivora, 21), we tested the influence of this sampling bias on uncertainty about ancestral states using stochastic character mapping of host orders and ecozones onto trees (35). We wrote a custom R function to resample tips from the phylogenetic tree and perform stochastic character mapping on the pruned tree using the packages `ape` and `phytools` (36, 37) assuming an equal-rates model. The function ran 100 mapping simulations on each pruned tree and calculated the probability that Chiroptera and Palearctic were the inferred host order and ecozone at the node uniting eubartonellae. These states were chosen based on initial reconstructions from BEAST indicating them as ancestral traits. We performed this simulation using three resampling schemes: equalizing the number of tips associated with bats and rodents (n = 87), equalizing tips associated with bats, rodents, and artiodactyls (n = 32), and equalizing tips associated with bats, rodents, artiodactyls, and carnivores (n = 21). Resampling schemes were run with 100 resampling iterations on the MCC tree and 10 resampling steps on 10 randomly sampled posterior trees. We summarized the resulting probability distributions by the mean and interquartile range (SI Appendix, Table S11). We further assessed the nature of transitions between hosts and ecozones by performing additional stochastic character mapping simulations on posterior trees followed by network analysis of state transitions. Host orders and ecozones were simulated with `phytools` over 1000 posterior sampled trees with an equal-rates model. The number of state transitions were then summarized over all 1000 simulations by the median and 95% credible intervals, ignoring state transitions with a median of zero (SI Appendix, Table S12). Separate host order and ecozone networks were then built from these median transitions, and node-level properties including degree, out-degree, and betweenness centrality were calculated using the R package `igraph` (38).

## Results

### Phylogeny and age estimation of *Bartonella* genus

Using molecular data from nine genetic loci sequenced from 331 *Bartonella* strains (SI Appendix, Table S1), we produced a well-supported Bayesian phylogeny (Fig. 1; SI Appendix, Fig. S8) that confirmed monophyletic clades of *Bartonella* species identified in past studies (19). These included a clade containing rodent-associated *B. elizabethae, B. grahamii, B. tribocorum*, and *B. rattimassiliensis* (clade H); a clade containing cat-associated *B. henselae* and *B. koehlerae* (clade F), *B. quintana*, and *B. washoensis* (clade E); and all three *B. vinsonii* subspecies (clade K). However, our approach has substantially altered the order of the deep branches within the phylogeny, including the delineation of five distinct *Bartonella* clades restricted to bats. Additional details regarding revisions to the *Bartonella* tree topology and clock rate estimates for sequenced loci can be found in SI Appendix.

Beyond a revised phylogeny, our approach demonstrated that bartonellae are ancient and supports the hypothesis that the genus diversified with mammals. We confirm that the genus first evolved as a symbiont of arthropods, represented by the species *B. apis, B. tamiae* and the ant symbiont *Candidatus* Tokpelaia hoelldoblerii, before transitioning to a parasitic lifestyle in mammals. These mammal-infecting eubartonellae (excluding *B. apis* and *B. tamiae*) began diversifying 62 million years ago (mya; 95% HPD: 40-90), near the Cretaceous-Paleogene boundary 66 mya (Fig. 1; SI Appendix, Fig. S6). Many crown metatherian and eutherian clades began diversifying around this time (21), including the diverse placental orders Chiroptera, Artiodactyla, Carnivora, Rodentia, and Primates, suggesting that *Bartonella* diversification is tightly linked with the radiation of its mammalian hosts during the Paleogene. Estimates of divergence dates using alternative substitution, tree, and clock models placed the origin of mammal-infecting eubartonellae between 57-70 mya (SI Appendix, Table S4).

### Diversification of bartonellae with hosts

Following the hypothesis that the *Bartonella* genus radiated with their mammal hosts, we performed tip-association tests to analyze the clustering of host taxonomic traits and geographic origin along the tips of the tree. Simulations using 1000 posterior sampled trees showed significant clustering of host orders and geographic ecozones across the phylogeny according to association indices (AI) and parsimony scores (PS). Observed distributions for both measures did not overlap their respective null distributions based on random associations of traits to tips (SI Appendix, Table S10). Host orders had smaller values for AI and PS than geographic origin, indicating a stronger phylogeny-trait association with host taxonomy than geographic origin. This phylogeny-trait association with host taxonomy is illustrated in Fig. 1A through strong support for monophyletic groups associated with host orders.

We clarified this association with host taxonomy by describing 15 *Bartonella* clades (SI Appendix, Tables S5 and S6) predominantly associated with marsupials (B), ruminants (C), carnivores (F), rodents (E, H, I, J, K, M, O), and bats (Fig. 1A). We then compared divergence dates of each *Bartonella* clade with divergence dates of the associated hosts within each clade (SI Appendix, Table S7) collated from TimeTree (32). We found a strong correlation between *Bartonella* and host clade divergence times (R^2^ = 0.72, F = 36.4, P < 0.0001). However, most (13/15) *Bartonella* clades were younger than their associated host clades; on average, the age of *Bartonella* clades was 76% that of their associated host clades (Fig. 2).

**Fig. 2.**
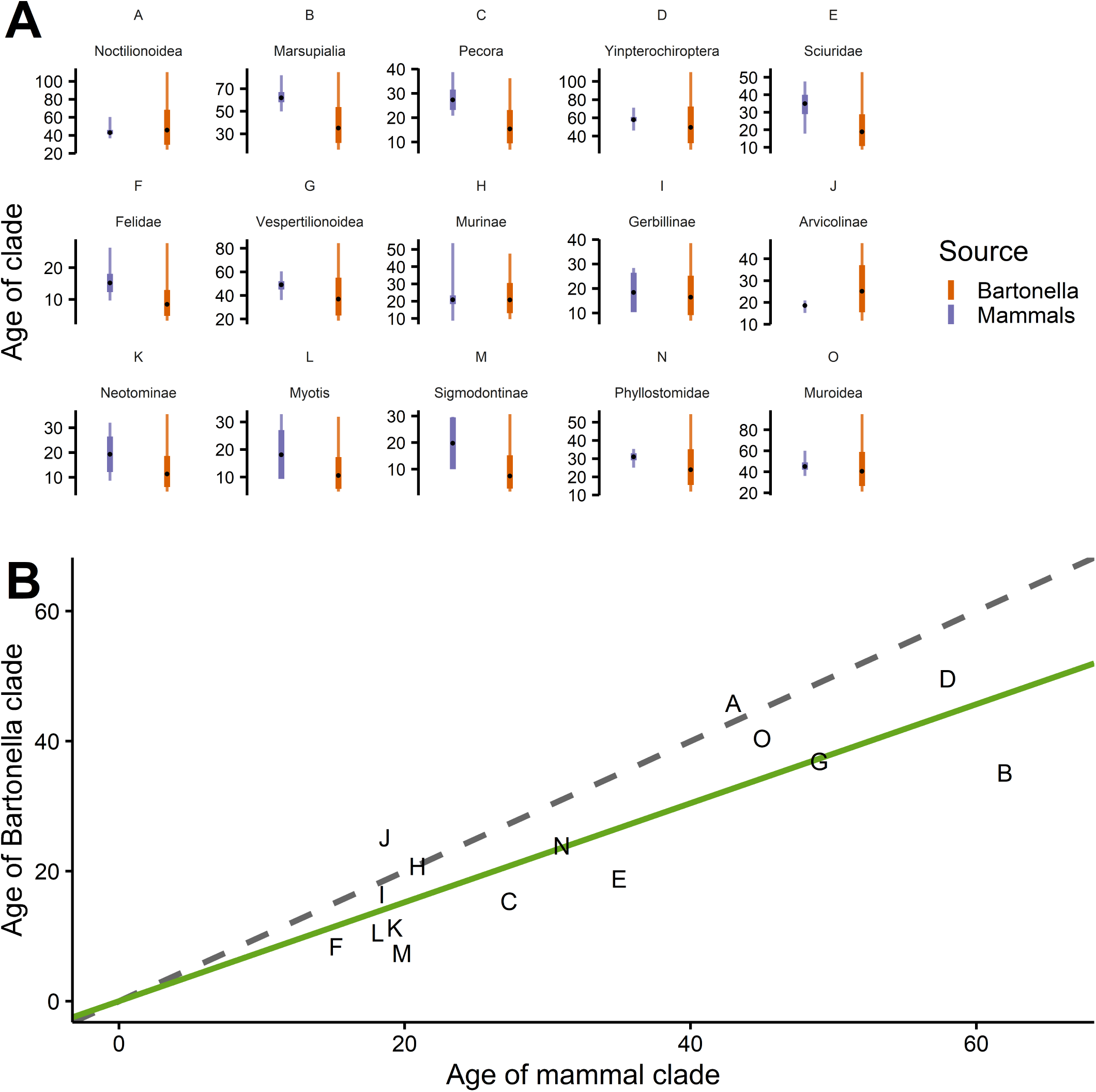
Comparison of divergence dates between *Bartonella* clades and associated host mammal clades. (A) Divergence times and intervals for *Bartonella* (in orange) and host clades (in purple). Black points show the median estimates and thin bars show divergence date ranges. Thick bars for mammal clades are the 95% confidence intervals estimated from TimeTree and the same bars for *Bartonella* clades are the 95% HPD intervals. (B) Correlation of median divergence dates between host and *Bartonella* clades, with clade identifiers shown as points. The solid green line indicates the best linear fit through the points and the dashed grey line shows the 1:1 line if host and *Bartonella* divergence dates were equal.

The Bayesian tree used in these analyses was similar to a maximum likelihood (ML) tree produced from concatenated sequences of all nine loci, with only minor differences in topology for some internal branches and external branches with low bootstrap support (SI Appendix, Fig. S7). Tip-association tests using the ML tree showed similar results to the Bayesian tree (SI Appendix, Table S10). Using confidence intervals for host clade divergence dates provided from TimeTree as calibration dates on the ML tree within the RelTime relative rate framework (39), we estimated the origin of mammal-infecting eubartonellae at 66.3 mya (95% CI: 63.5-69.1). This separate analysis validates the Bayesian relaxed clock estimate (SI Appendix, Table S5) and further supports the inference that *Bartonella* began diversifying with mammals near the Cretaceous-Paleogene boundary.

### Influence of host groups and geography on *Bartonella* evolution

Bats appear to be highly influential in the diversification and spread of *Bartonella* geographically and to other host orders. Bat-associated clades (A, D, G, L, N) are broadly distributed across the tree and form external branches to clades associated with other mammalian orders (Fig. 1A). This contrasts with clades associated with marsupials, ruminants, carnivores, and rodents, which are less dispersed on the tree and stem from more internal branches. Based on ancestral state analysis using host orders as states, bats were inferred to be the ancestral host of all mammal-infecting eubartonellae with a posterior probability of 0.99. Due to the large number of bat-associated strains in the database (n = 160), this inference of the ancestral host may have been biased towards bats. Yet in all resampling scenarios, the median posterior probability that bats are the ancestral hosts of mammal-infecting eubartonellae exceeded 0.9 (SI Appendix, Table S11). In further support of this inference, the diversification of mammal-infecting *Bartonella* started almost exactly when bats began their evolutionary radiation around 62 mya (95% CI: 59-64, range: 51.9-74.9) according to compiled studies from TimeTree (32).

In addition to ancestral host associations, we also inferred the ancestral biogeography of *Bartonella* clades and where host transitions may have occurred. We performed ancestral state reconstruction of ecozones based on the current geographical distribution of the host of each *Bartonella* strain. The geographical origin of eubartonellae was inferred to be in the Palearctic (Fig. 1B) with a posterior probability of 0.99. This fits with the classification of bats within the clade Laurasiatheria and previous reconstructions of chiropteran biogeography which found that extant bats may have originated in Eurasia (40). However, the inference of the geographic origin of eubartonellae is less certain when host sampling bias was accounted for in the stochastic character mapping analysis. The median posterior probability for a Palearctic origin of eubartonellae ranged from 0.63 to 0.77 across all resampling scenarios (SI Appendix, Table S11). Regardless of the exact geographical origin, it is probable that bats have been influential in the ancient geographic spread of *Bartonella* infections (Fig. 1).

We explored the influence of particular hosts on the spread of *Bartonella* among mammalian orders and across ecozones using stochastic character mapping and network analysis. After mapping the number of host and ecozone transitions across 1000 posterior sampled trees, we built a network consisting of host and ecozones as nodes and the median number of transitions between nodes as edges (Fig. 3; SI Appendix, Table S12). In general, the ecozone network was more highly connected than the host network (Fig. 3). The higher number of connections in the ecozone network corresponds with the results of the tip-association tests (SI Appendix, Table S10), which showed that clustering of traits was stronger for host taxonomy than geographic origin. That is, the high frequency of transitions between ecozones leads to lower levels of geographical clustering on the tree.

**Fig. 3.**
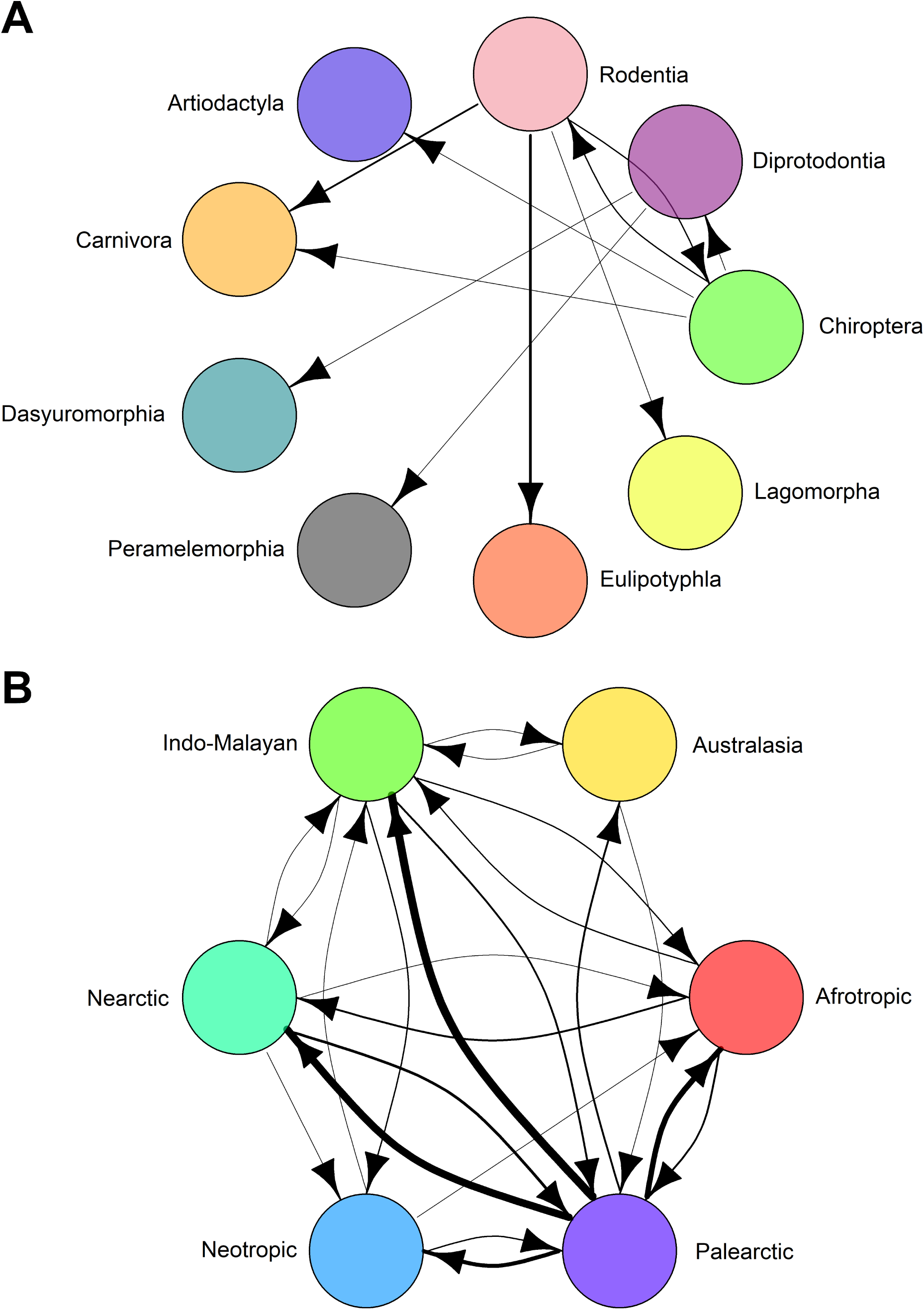
Transition network for (A) host orders and (B) ecozones across the *Bartonella* phylogeny. Edges connecting nodes are the median number of transitions between host and ecozone states based on stochastic character mapping on 1000 posterior sampled trees. Edge widths are proportional to the median number of transitions. Edges with a median of zero transitions are not shown. Transitions between the outgroup (*Brucella abortus*) and between mammalian orders and arthropods have been removed for clarity. All transition counts with a median above zero are shown in SI Appendix, Table S9.

Examining the network properties of the nodes, we find that certain host orders are influential in the spread of *Bartonella* among host orders (SI Appendix, Table S13). In particular, we considered degree (the number of edges connected to a node), out-degree (the number of edges originating from a node), and betweenness (the number of shortest paths that connect any two nodes in the network that pass through the node in question) because these measures describe how each node serves as a source of *Bartonella* to other nodes. Bats and rodents were a source to other mammalian orders (Fig. 3A), with the highest degree and out-degree of all host orders and high betweenness (SI Appendix, Table S13). Transitions between ecozones show that the historical movement of *Bartonella* by hosts led to the present global distribution of these bacteria (Fig. 1B) through bidirectional exchange (Fig. 3B). Palearctic and Indo-Malayan ecozones showed the highest degree, out-degree, and betweenness. Thus, these two regions may have played an important role as geographic hubs for *Bartonella* diversification and movement of hosts to other ecozones (Fig. 1B; SI Appendix, Fig. S8B).

## Discussion

*Bartonella* is a broadly distributed bacterial genus associated with mammals and arthropod vectors globally. Patterns of host-specificity and phylogenetic diversity in this genus reflect general trends in other zoonotic pathogens. Thus, *Bartonella* serves as a model system for understanding the evolution and ecology of zoonotic agents. Specifically, this system could inform theory about how agents adapt to and diversify in hosts over time and the ecological conditions that lead to accidental infections and host-switching. Using a multi-faceted analytical approach, this study answered several key questions about the evolution of *Bartonella* bacteria. First, we found that the *Bartonella* genus began diversifying with mammals around the Cretaceous-Paleogene boundary. Our novel approach used a strong relaxed clock prior on the 16S rRNA locus based on substitution rates observed in bacterial symbionts of arthropods (17) while accounting for rate variation at eight other genetic loci to yield a highly supported phylogenetic tree with estimated divergence dates. Second, we showed that *Bartonella* clades diversified along with their mammalian hosts. Ancestral state reconstruction on the phylogenetic tree showed that *Bartonella* lineages tend to cluster by host taxonomic orders and this clustering was found to be significantly higher than random expectations using tip-association tests. Additionally, we found a significant correlation between the divergence times of 15 *Bartonella* clades and their associated host clades. A separate time tree estimation approach calibrated using these host divergence dates confirmed the dating of eubartonellae diversification. The use of ancestral state reconstruction or stochastic character mapping of host traits paired with network analysis is a nascent approach in the study of infectious agents that can provide additional insights from phylogenies (41–43). These analyses demonstrated that bats have been key to both the origin and spread of *Bartonella* among other mammals and geographic regions, while rodents were responsible for additional spread. This work elucidates key aspects of the ecology and evolution of *Bartonella*, yet there are several avenues of research to be explored in future studies.

One necessity is to thoroughly catalog *Bartonella* diversity. While description of *Bartonella* species was slow through the 20^th^ century, the advent of genetic sequencing has brought about an explosion of *Bartonella* diversity with over 40 named and likely many other unnamed species. Our phylogenetic analysis used the most comprehensive sequence database to date, including broad taxon sampling of *Bartonella* strains characterized from 10 mammalian orders. These data, along with a relaxed clock approach, have reshaped the *Bartonella* phylogeny, defining five new clades of bat-associated *Bartonella* strains and reorganizing the relationships of deeply branching clades. Attempts to culture and characterize novel *Bartonella* strains from undersampled mammalian orders or other potential vertebrate hosts (e.g., birds (44)) are needed to further improve taxon sampling. This continued work will undoubtedly reshape the *Bartonella* tree further and may lead to new hypotheses about ancient associations with hosts.

Our results also provide context to the biological changes that are associated with the shift of *Bartonella* bacteria from an arthropod symbiont to a mammal parasite. Our phylogeny reaffirms work demonstrating this shift (16, 18) and provides an estimated time for when it occurred, suggesting that an existing bacterial population colonized a new niche in mammals shortly after their emergence as potential hosts. Some of the molecular machinery that could have facilitated this colonization was already present in arthropod-associated *Bartonella* lineages and other Rhizobiales bacteria (16). The majority of virulence factors important for host interaction or establishment of intracellular infection are shared across Bartonellaceae, suggesting some latent potential for infecting vertebrates even in arthropod-associated lineages. However, the evolutionary radiation of eubartonellae is associated with a number of other important molecular innovations, including the loss of flagella and acquisition of *trw* and *virB* type IV secretion systems (T4SS) (16, 19, 45, 46). Secretion systems have only been detected and characterized in a few *Bartonella* species across the phylogeny, so our revision of *Bartonella* tree topology highlights a need for future work regarding the machinery (e.g., flagella, T4SS) shared between bat-associated lineages and their relatives.

Given that current mammal-associated bartonellae are vectored by blood-feeding arthropods and ancestral bartonellae were likely arthropod symbionts, it is probable that early adaptation to blood-feeding arthropods facilitated the colonization of the mammalian bloodstream. Hematophagous arthropods frequently harbor endosymbionts to cope with their nutritionally deficient diet (47), so ancient (and possibly some extant) bartonellae may have had beneficial relationships with arthropod hosts. The switch from symbiont to mammal parasite could then have occurred early in the evolution of mammals. There is evidence that ancestors of extant mammalian ectoparasites implicated as *Bartonella* vectors (48–50) were already present by the end of the Cretaceous, including sand flies (51), fleas (1), sucking lice (52), bed bugs (2), and hippoboscoid flies (53). Based on available evidence, the colonization of mammals by *Bartonella* bacteria may have occurred via hematophagous vector, possibly parasitizing early bats. An ancestral relationship with bats is supported by recent detection of *B. tamiae* in bat flies and bat spleens (54, 55), suggesting that this species can opportunistically colonize bats from arthropods even today. The initial transmission may have occurred through contamination of skin with arthropod feces containing bacteria, direct consumption of an infected arthropod, or some other unknown route (4). Once inside the host, the existing ability of bartonellae to invade host cells may have led to proliferation of bacteria in the blood. Additional studies that isolate *Bartonella* lineages in arthropods and confirm potential transmission routes between mammal hosts and arthropod vectors will clarify the evolution of host-vector-*Bartonella* relationships.

As apparent in Figs. 1 and 3, the evolutionary history of *Bartonella* has involved several host-switching events. Thus, calibrating divergence dates by relying on codivergence between host taxa would poorly reflect this history. Instead we initially avoided a calibration approach in favor of using a relaxed clock prior, then validated estimated divergence dates based on 15 radiation events within particular bat, rodent, ruminant, and marsupial host taxa. The *Bartonella* divergence dates correlate strongly with the host divergence dates, although with a widespread delay in the colonization of *Bartonella* within a clade (Fig. 2). While it is possible that this delay in *Bartonella* colonization is associated with the divergence date estimation approach and bacteria diverged immediately along with their hosts, we suspect the delay reflects some biological reality. According to Manter’s rules (56, 57), parasites evolve more slowly than their hosts due to the relatively uniform environments they experience within a host. This slow evolution may help to explain rampant *Bartonella* host-switching between related hosts in the tree, since from a parasite’s perspective the intracellular environments of phylogenetically similar hosts are unlikely to have significantly changed. Despite these inherent delays, the clustering of *Bartonella* strains with host orders and particular clades within those orders along with the correlation of divergence times strongly suggest a shared evolutionary history between *Bartonella* strains and their hosts, although a more complicated one than simple cospeciation.

Beyond patterns of codiversification, it is clear from this study that *Bartonella* evolution has been shaped by certain hosts, particularly rodents and bats. As the two most speciose groups of mammals, they could be expected to host diverse parasites according to Eichler’s rule (58), which predicts positive covariance between host and parasite diversity. While more studies will need to be done to explicitly test patterns of host and *Bartonella* diversity while accounting for sampling biases, it is clear from the network analysis that rodents and bats are important sources of bartonellae to other hosts (Fig. 3). As abundant taxa within ecosystems, rodents and bats could act as targets for both generalist and specialist ectoparasites. While endemic *Bartonella* infections are likely maintained by transmission by specialist ectoparasite vectors, generalist vectors could target the most abundant species in the community (e.g., rodents or bats) and occasionally infest alternative hosts, resulting in opportunities for accidental *Bartonella* infections in phylogenetically distant hosts over evolutionary time (31). Reconstructing some of these ancient host-switching dynamics would require knowledge of ancestral ectoparasite associations and the interactions of hosts and their ectoparasites within communities.

Finally, bats were identified as the most probable ancestral host of eubartonellae in mammals even after accounting for sampling bias in the database. The fact that bats can fly would have hypothetically increased their dispersal ability during their early diversification. This is exemplified by numerous long-distance colonization events: from mainland Africa to Madagascar by seven different extant bat families, including the endemic Myzopodidae; from Australia to New Zealand by the family Mystacinidae; and from mainland North America to Hawaii by *Lasiurus cinereus* (59). The dispersal of bats to distant landmasses during the early diversification of extant mammals could have played a role in the importance of bats as sources of *Bartonella* infection to other hosts. We also note that bats appear to be highly tolerant of infections, especially of intracellular bacteria and viruses (60), showing few signs of disease and unique immune responses compared to other mammals (7, 61–63). Such patterns in extant bats may have ancient origins linked with their ability to fly (64) and thus bats may have been ideal hosts for the early colonization of mammals by arthropod-borne bartonellae.

The importance of bats in the evolutionary diversification of mammal parasites has been discussed by other authors working in distinct systems. One of these groups are the *Trypanosoma* parasites that include *T. cruzi*, the agent of Chagas disease. Observing the broad distribution of bat-associated clades in the growing diversity of trypanosomes, Hamilton and others hypothesized that bats may have been highly influential in the geographic spread of the *T. cruzi* clade and host-switching to other mammals (14). This ‘bat-seeding’ hypothesis has continued to gain support since it was proposed with the discovery of diverse lineages in the *T. cruzi* clade in bats globally (12, 41). Similar patterns have been noted in malarial parasites (Haemosporida), wherein the transition from sauropsids into mammals likely occurred only once, with bats being a possible bridge to other mammals (13, 65). In light of the results of this study and the patterns in other systems, we contend that the ‘bat-seeding’ hypothesis may apply more widely among mammalian parasites. Our approach using comprehensive phylogenetic analysis, estimation of divergence times, and ancestral reconstruction of host associations could be applied to understand the evolutionary radiation and host-switching patterns of these parasites, and potentially the role that bats have played in their diversification.

## Acknowledgements

We thank the members of the Webb laboratory for their suggestions on early drafts of this manuscript. The findings and conclusions in this report are those of the authors and do not necessarily represent the official position of CDC.

## Author contributions

C.M. and M.K. designed research; C.M., Y.B., and M.K. performed research; C.M. analyzed data; and C.M. and C.W. wrote the paper.

