## Supplemental Text for "Bats are key hosts in the radiation of mammal-associated *Bartonella* bacteria"

### Molecular data collection: further details

*DNA extraction and sequencing of bat-associated* Bartonella *strains*

Genomic DNA was extracted from 129 bat-associated *Bartonella* cultures using a simple heat extraction protocol (incubation at 95°C for 10 min) and diluted 1:10 in extraction buffer (Qiagen, Valencia, CA). Amplification of targeted genetic loci (Table S1) used published primers and protocols (1–4). Amplification of *groEL* was unsuccessful for many strains with the available primers (5), so this locus was not sequenced for any bat-associated strains and was only available from MLSA and genomic data. Positive PCR amplicons were purified using the Qiagen QIAquick PCR Purification Kit and sequenced in both directions with the same primers on an Applied Biosystems Model 3130 Genetic Analyzer (Applied Biosystems, Foster City, CA). Reads were then assembled in Lasergene v14 (DNASTAR, Madison, WI). Repeated amplification or sequencing was performed for some missing genes, but for 28 strains there was one or more sequence that could not be obtained: *ftsZ* (2), *nuoG* (2), *ribC* (22), or *rpoB* (3).

*Sequence alignment and data cleaning*

For all bat-associated and reference *Bartonella* strains, sequences from each genetic locus were aligned separately with MAFFT v7.187 (6). Ends of alignments and poorly aligned sites were trimmed with Gblocks v0.91b (7), and final alignments were manually checked for ambiguous base pairs and edited. The final alignment lengths and coverage across taxa are listed in Table S1, with an average of 78% coverage across the nine loci. We concatenated all loci using Phyutility v2.2 (8) to produce a full supermatrix of 8345 base pairs (including gap sites) for later analyses. Preliminary phylogenetic analysis of bat-associated strains determined that seven showed evidence of homologous recombination with another bat-associated strain (even after repeated amplification and sequencing) and one showed highly discordant phylogenetic positions across sequenced loci, so these strains were removed from the database.

*Molecular data validation*

Previous analyses have shown that the protein-coding loci (*ftsZ*, *gltA*, *groEL*, *nuoG*, *ribC*, *rpoB*) are under purifying selection with low ratios of non-synonymous to synonymous substitutions (1, 4). The 16S rRNA locus is known for being highly conservative within a bacterial genus (9, 10). As a spacer sequence, ITS is unlikely to be under selection. We examined GC content across the full alignment for all 332 taxa using DAMBE v7.0.48 (11). The eubartonellae clade and *B. tamiae* exhibited a stationary GC content distribution between 0.38-0.48 while *B. apis*, *Candidatus* Tokpelaia hoelldoblerii, and *Brucella abortus* had progressively higher GC content values (Fig. S1). Previous studies have shown that the similarity in GC content between *B. tamiae* and eubartonellae can affect phylogenetic results. Specifically nucleotide alignments show that *B. tamiae* is a sister taxon to eubartonellae while protein alignments or nucleotide alignments without the third codon position show that *B. tamiae* is a sister taxon to *B. apis* (12, 13). Since this inference was focused primarily on the eubartonellae clade and not on its putative sister taxa in arthropods, we determined that the stationary GC content distribution for eubartonellae was acceptable for phylogenetic analysis and required no correction.

To confirm the absence of homologous recombination within taxa in the database, we generated a network phylogeny in SplitsTree v4.14.8 (14) using the concatenated alignment and the Neighbor-Net method (15) on uncorrected pairwise distances . The network phylogeny showed a moderately tree-like structure (Fig. S2) with parallelograms connecting closely related taxa and basal splits indicative of shared evolutionary history. A pairwise homoplasy (PHI) test (16) for recombination implemented in SplitsTree found no statistically significant evidence for recombination ($P = 1$) for the concatenated alignment or each locus separately (Table S2).

Separate loci were tested for the presence of nucleotide substitution saturation by plotting uncorrected versus adjusted distances (Tamura-Nei model (17)) and transitions/transversions versus adjusted distances using DAMBE and the R package `ape` (18–20). Adjusted distances did not show substantial saturation, exhibiting a strongly linear relationship with only slightly asymptotic behavior at the farthest distances (Fig. S3). Transitions and transversions fell along a straight line (Fig. S4) and transitions largely outnumbered transversions for all loci except ITS (11), indicating no substantial evidence of saturation. The absence of significant saturation was confirmed for all loci (Table S3) in DAMBE using the test developed by Xia et al. (21). Based on all the tests above, we determined that these molecular loci would be appropriate for phylogenetic analysis and accurate estimation of divergence times.

**Phylogenetic analysis: further details**

*Phylogenetic model selection*

The best sequence evolution model was chosen according to the Akaike information criterion (AIC) using jModelTest v2.1.6 (22) via the CyberInfrastructure for Phylogenetic RESearch (CIPRES) Science Gateway portal v3.3 (23). The generalized time-reversible model with a proportion of invariant sites and gamma rate variation across sites (GTR+I+G) was chosen for all loci except *ssrA*, which best fit the Tamura-Nei model (TN+I+G) (Table S1). Since substitution models had little effect on the tree topology, we chose to analyze all loci using the GTR+I+G for consistency and to correspond with the maximum likelihood analysis, which used a GTR+I+G model. A maximum likelihood (ML) tree was generated from the concatenated alignment of nine loci using RAxML v8.2.12 (24) on CIPRES with 1000 bootstrap iterations to estimate node support. The ML tree was used to compare topologies with the Bayesian tree and for tip-association tests.

*Phylogenetic model priors and run settings*

The prior distributions for substitution rate and speciation model parameters are listed in Table S4. The prior for all ancestral state transitions was a gamma distribution with shape and scale parameters set to one, and the prior for the mean rate of order and ecozone transitions were set to the CTMC approximate reference prior (25). We ran three chains in BEAST using the model settings above for the final analysis. The chains were run for $2x10^8$ iterations, sampling parameters every $2x10^4$ iterations. We inspected posterior distributions for all model parameters to assess convergence, mixing, and high effective sample sizes (ESS > 200) using Tracer v1.7.1 (26). We chose the maximum clade credibility (MCC) tree from the posterior tree iterations after burn-in using TreeAnnotator (26) and the tree with the highest MCC score was used for all subsequent analyses. The final tree was visualized and edited in FigTree v1.4.4. Molecular clocks for the nine genetic loci were summarized by the median and highest posterior density (HPD) of their distributions. The divergence date of the most recent common ancestor of mammal-infecting *Bartonella* (eubartonellae, excluding *B. tamiae* and *B. apis*) was summarized from the MCC tree by the median and HPD.

*Testing alternative models in BEAST*

To increase confidence in the robustness of our conclusions with respect to phylogenetic model choice, we performed additional runs in BEAST using alternative models and subsets of sequence data. The amount of data and the complexity of models led to long computational runtimes (up to 7 days) that reached the limit permitted on CIPRES (23). For this reason, we did not pursue a formal model selection approach through estimation of marginal likelihoods (27, 28) and instead chose to run a non-exhaustive series of models using combinations of alternative model settings to assess the combined effects on the topology and divergence times on the resulting tree.

For models that used a TN+I+G model for *ssrA*, the two prior distributions for the kappa priors were chosen to be lognormal with log mean of one and a log standard deviation of 1.25 with an initial value of 2. For models that used an uncorrelated relaxed clock model with a lognormal distribution of clock rates along branches, the means for each locus were set as for the exponential distribution detailed in the main text. An additional prior was set for the standard deviation of the lognormally distributed clock rates using an exponential distribution with a mean of 0.33. All model combinations were run until parameters converged to stationary distributions as determined through visual inspection in Tracer v1.7.1 (26). Burn-in iterations were removed and the maximum clade credibility (MCC) tree was selected using TreeAnnotator (26). We then compared the topology and divergence dates (particularly the estimated divergence date of eubartonellae) of the MCC trees.

Regardless of substitution, codon partitioning, clock, or tree models, we found only limited variation in the topology of the tree across all runs with no major changes in the position of large clades that would influence the results or conclusions in the main text. The divergence dates of eubartonellae (Table S4) and *a posteriori* defined clades (Table S8) varied little across runs, indicating that the molecular data, taxon sampling, and choice of prior on the 16S rRNA clock were more important to phylogenetic inference than any other model settings. The only major differences observed in the topology and divergence dates of the tree were observed when a strict clock was used. These runs predicted a younger divergence date for eubartonellae (~57 mya) and showed a different arrangement of clades A-C and the clades that contain *B. bacilliformis*, *B. rochalimae*, and *B. clarridgeiae*. All runs using strict molecular clocks had much lower likelihoods than runs using relaxed clocks, so the use of a strict clock was rejected. As long as variation in clock rates were allowed to be uncorrelated across the branches of the tree, the topology and divergence dates on the tree were stable.

We note that the exclusion of ITS sequences had little effect on tree topology and divergence dates, so this locus may have had limited phylogenetic signal. Nevertheless, we retained this locus for the final run used in the main text. Since codon partitioning and the choice of relaxed clock models had little influence on the trees, we chose not to use codon partitioning and to use exponential distributions for the uncorrelated relaxed clocks rather than lognormal distributions for the final runs in the main text to reduce the number of independent parameters that needed to be estimated.

### *Bartonella* lineages associated with arthropods: further details

Several *Bartonella* lineages in the database were labeled as being associated primarily with arthropods for the ancestral state reconstruction analysis. *B. apis* was originally isolated from western honey bees (*Apis mellifera*) and has not been associated with any mammalian hosts (29). While several strains of *B. apis* have been characterized from honey bees in North America and Europe (29), we chose to associate this species with the Palearctic ecozone to reflect the hypothesized historical distribution of domesticated *Apis mellifera* in northern Africa or the Middle East (30). *B. tamiae* was originally isolated from humans in Thailand (31), this likely represents an accidental association. Genetic sequences identified as *B. tamiae* or closely related to this species have been obtained from several arthropod species including bat flies and bat ticks (32, 33) and chigger mites collected from rodents (34). Given its basal position relative to the mammal-associated eubartonellae clade and closer affinities with *B. apis* (29), we chosen to associate *B. tamiae* primarily with arthropods for this analysis.

We labeled *B. bacilliformis* and its relatives *B. ancashensis* and *Candidatus* B. rondoniensis as being associated with arthropods instead of a particular mammalian order because a reservoir host has not been conclusively determined for these species. *Bartonella bacilliformis* causes severe morbidity and mortality in humans, and prevalence is generally low in human populations, so humans are unlikely to be the reservoir host (35). Furthermore, repeated attempts to isolate *B. bacilliformis* from alternative plant or animal reservoirs have been unsuccessful (35). Despite the uncertainty about the reservoir host, *B. bacilliformis* is known to be vectored by *Lutzomyia* spp. sandflies (36–38). A recent study also reported the presence of *B. bacilliformis* in ticks collected from tapirs and peccaries in Peru (39). The phylogenetically related *Candidatus* B. rondoniensis was also described from the assassin bug *Eratyrus mucronatus* in French Guiana (40). While the host or vector of *B. ancashensis* is unknown (41) it is part of a clade that includes *B. bacilliformis* and *Candidatus* B. rondoniensis. Given the uncertainty of the mammalian hosts for this *Bartonella* clade, we chose to associate this group primarily with arthropods since it appears to be the ancestral trait (42). Future work that conclusively determines the mammalian hosts of *B. bacilliformis* and its allies is clearly necessary and could improve the inference of ancestral hosts for *Bartonella* lineages.

Finally, the host origin of *B. senegalensis* is unclear since it was isolated from the soft tick *Ornithodoros sonrai* in Senegal (43). Although the ticks were found in rodent burrows, the presence of the bacteria was not confirmed in any mammals, so we chose to associate this bacteria with arthropods. Similar to the clade that includes *B. bacilliformis*, future studies involving this bacteria and its host associations will improve our knowledge of evolution within the *Bartonella* genus.

We confirmed that none of these choices had an effect on the results by repeating stochastic character mapping using alternative assignments of traits to these tips. Bats were always inferred to be the ancestral hosts of eubartonellae. Thus, we chose to retain these trait assignments for the analysis in the main text.

**Revision of *Bartonella* tree topology: further details**

In contrast with past phylogenetic analyses of the *Bartonella* genus that used only maximum likelihood analysis of concatenated genes (44–48), we showed that neither *B. bacilliformis* nor *B. australis* are the most deeply branching lineages in the genus. Instead we found that *B. bacilliformis* and its allies *B. ancashensis* (41) and *Candidatus* B. rondoniensis (40), constituting the clade previously named "Lineage 1", are in fact most closely related to ruminant-associated *Bartonella* species including *B. bovis*, *B. schoenbuchensis*, and others (Fig. 1; Fig. S8); a finding supported by Wagner and Dehio (45). This ruminant clade was named clade C in our analysis and "Lineage 2" in other studies (44–48). The group of species including *B. rochalimae, B. clarridgiae*, and allies, previously named "Lineage 3", was found to be distantly related to a clade containing kangaroo-associated *B. australis* (49) and other *Candidatus* strains from marsupials (50, 51), and two lineages associated with bats, one from Africa (1) and one from Europe (52). These clades ("Lineages 1-3") were all found to be part of a strongly supported monophyletic clade (posterior probability, PP = 1) that includes the deeply branching sister group clade A associated with neotropical bats (Fig. 1). The bat-associated and marsupial-associated clades could potentially be elevated to the level of lineages equal to the others. Alternatively, unification of these lineages into a monophyletic clade would suggest a redefinition of lineages into subclades.

Broad taxon sampling also expanded "Lineage 4", a well-supported clade (PP = 1) that contains all other *Bartonella* species separate from "Lineages 1-3" and most of the diversity in the genus (Fig. 1). Specifically, we discovered four new bat-associated clades (D, G, L, N) within this lineage, with clade D as the sister group to all other "Lineage 4" clades and clade G as the sister group to a large clade of predominantly rodent-associated *Bartonella* species (enclosed within clade O). Clade L, containing strains from North American and European vespertilionid bats (52–55) and *Candidatus* B. mayotimonensis (56), along with clade N associated with neotropical bats, are contained within a clade that includes the *B. vinsonii* species complex associated primarily with rodents (57–63). We also recovered a monophyletic clade (PP = 1) that unites rodent-associated *B. birtlesii*, *B. doshiae*, and *B. taylorii*, similar to a previous MLSA study (4). Subdivisions within lineage 4 could be based on radiations within distinct mammalian groups, as we have done (Fig. 1A; Tables S5-S6). This revision of the *Bartonella* tree through increased taxon sampling and characterization of bat-associated strains illustrates the diversity in this genus that remains uncharacterized.

**Estimated clock rates for *Bartonella* genetic markers: further details**

To verify that the molecular clock approach could capture variation across loci using a single strong prior distribution on the 16S rRNA gene, we analyzed clock rates for each of the nine loci. Clock rates predictably varied by gene function (Table S9). The 16S locus had a very low median clock rate at $5.2 \times 10^{-10}$ nucleotide substitutions site$^{-1}$ year$^{-1}$ (95% HPD: 3.4-$7.1 \times 10^{-10}$) across branches. As this locus codes for a functional RNA with a conserved 3D structure, this low rate was deemed reasonable and was very close to previous estimates of 16S rRNA divergence of 1-2% per 50 million years in *E. coli* and *Buchnera* symbionts of aphids (64, 65). Protein-coding loci and the functional transfer-messenger RNA locus *ssrA* had branch rates five to nine times higher than 16S rRNA while ITS had rates 22 times higher than 16S rRNA (Table S9).

**Biogeographic patterns of select *Bartonella* clades: further details**

In support of our hypothesis that *Bartonella* clades co-diverge with their mammalian hosts, there were several instances of deep separations of host-associated *Bartonella* strains that are most compatible with an ancient origin of the *Bartonella* genus. Inoue *et al.* discovered phylogenetically distinct clades of *B. washoensis* infecting ground squirrels in North America and Asia (66), a result we replicated in our tree within clade E (Fig. 1; Fig. S8). The squirrels harboring these bacteria are from two separate genera, *Spermophilus* from Eurasia and *Urocitellus* from North America, that diverged 7.8 mya according to studies published on TimeTree (67). Therefore, it is unlikely squirrels from these two genera have been in recent close contact that could lead to *Bartonella* transmission and the divergence observed in the *Bartonella* clades reflects their independent evolution in isolated hosts. Similar patterns were seen in *Bartonella* clades infecting bats. One involves the separation of two clades within bat-associated clade L (Fig. 1). One clade within this group (Fig. S8) contains strains from vespertilionid bats in Europe (52, 54) and the other clade contains a strain from North American bats (53) and an agent of human endocarditis, *Candidatus* B. mayotimonensis (56). *Myotis* and *Eptesicus* spp. bats in North America diverged from their congeners in Eurasia 16.2 and 15.3 mya respectively according to TimeTree. Within the large clade D harbored by Old World bats (Fig. 1; Fig. S8) there are two *Bartonella* strains infecting *Hipposideros* spp. bats, *H. larvatus* from Thailand (2) and *H. vittatus* from Kenya (68). While *Hipposideros* species have repeatedly moved between Africa and Asia according to phylogenetic analysis (69), these two species have been separated for 34 million years. These divergence times between geographically isolated hosts are reflected in the estimated times for their *Bartonella* divergence times: 5.2 mya for *B. washoensis*-like strains in ground squirrels, 10.5 mya for *Candidatus* B. mayotimonensis-like strains in vespertilionid bats, and 27.6 mya for the two *Hipposideros*-associated strains. These results provide confidence in the molecular clock approach and an ancient diversification of the *Bartonella* genus, however more work is clearly needed to reconstruct historical biogeographical patterns of bartonellae and their hosts.

**Network analysis of *Bartonella* host and ecozone transitions: further details**

Based on our stochastic character mapping of host orders and ecozones onto 1000 posterior sampled trees, we built separate networks using host orders and ecozones as nodes and the median number of transitions as edges. Considering only transitions within and between mammalian orders, the ecozone network had 22 non-zero median transitions between 6 nodes, resulting in a network density of 73% considering all possible directed transitions. The host order network showed only 10 non-zero transitions between 9 nodes, for a density of 14%. The ecozone network also had higher median counts of transitions than the order network, with up to 12 observed transitions between the Palearctic and

Indo-Malayan ecozones (Table S12). Rodents are a source of transitions to Carnivora, Eulipotyphla, and Lagomorpha, while bats are a source to Diprotodontia and other marsupials, Artiodactyla, and Carnivora. Many of these transitions are strongly supported within the MCC tree with posterior probability greater than 0.9 (Fig. 1A; Fig. S8A). Notable transitions from Rodentia include those to Carnivora at the ancestor to *B. rochalimae* and to *Bartonella* sp. JM-1 from *Martes melampus* within clade E (including *B. washoensis*); to Eulipotyphla at the ancestor to *B. florenciae*, to *Bartonella* sp. DB5-6 from *Sorex araneus* within clade J (including *B. birtlesii*, *B. doshiae*, and *B. taylorii*), and to *B. tribocorum* and *B. queenslandensis* strains from shrews within clade H; and to Lagomorpha at the ancestor to *B. alsatica*. Well-supported transitions from Chiroptera include to Diprotodontia and other marsupials at the ancestor to clade B (including *B. australis*). Rodents and bats showed an equal number of transitions between each other (Fig. 3A), however the sources of these transitions are equivocal with lower posterior probabilities for the ancestral host (Fig. 1A).

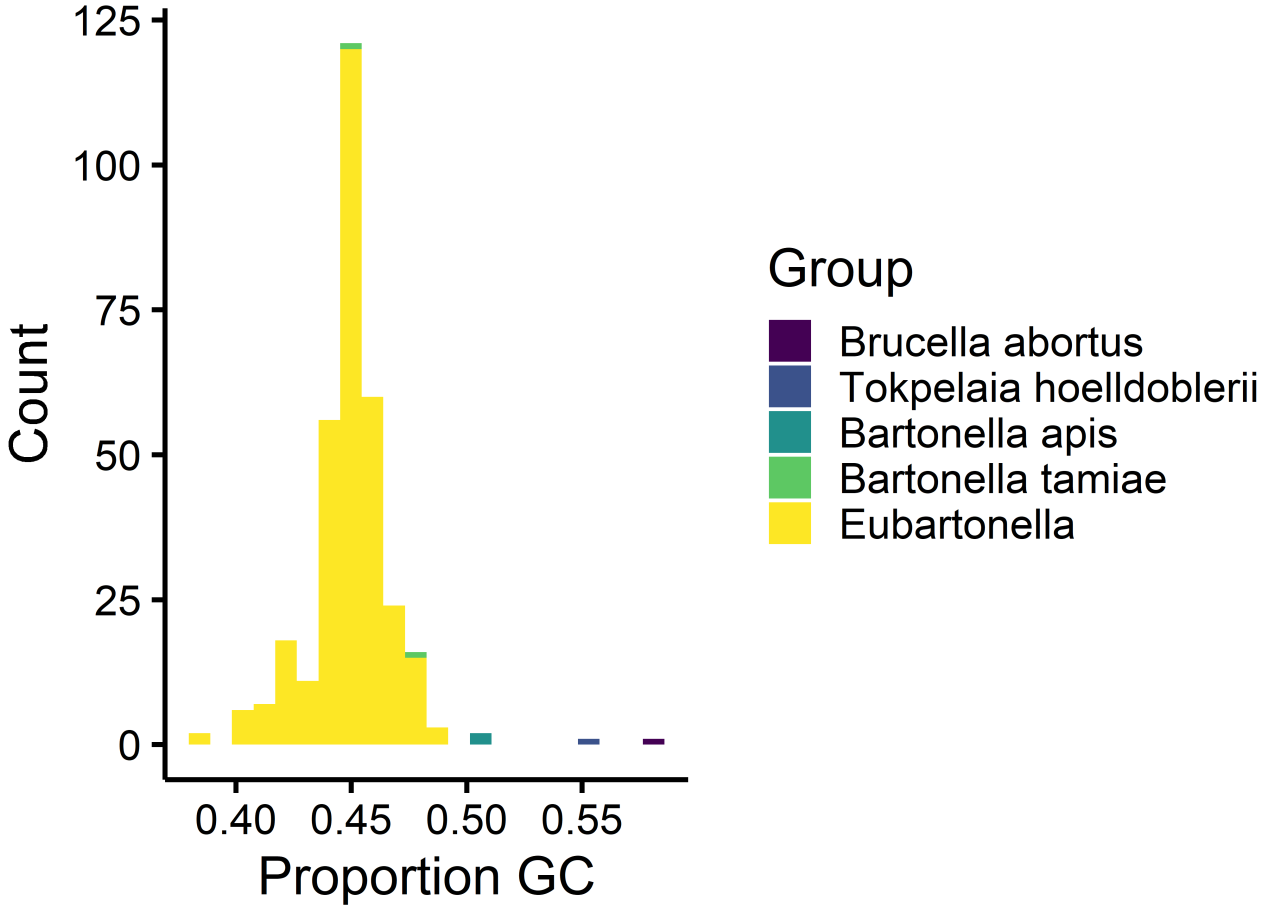


**Fig. S1.** Histogram of GC content across taxa. Nucleotide content was calculated across the concatenated alignment of nine loci for all 332 taxa.

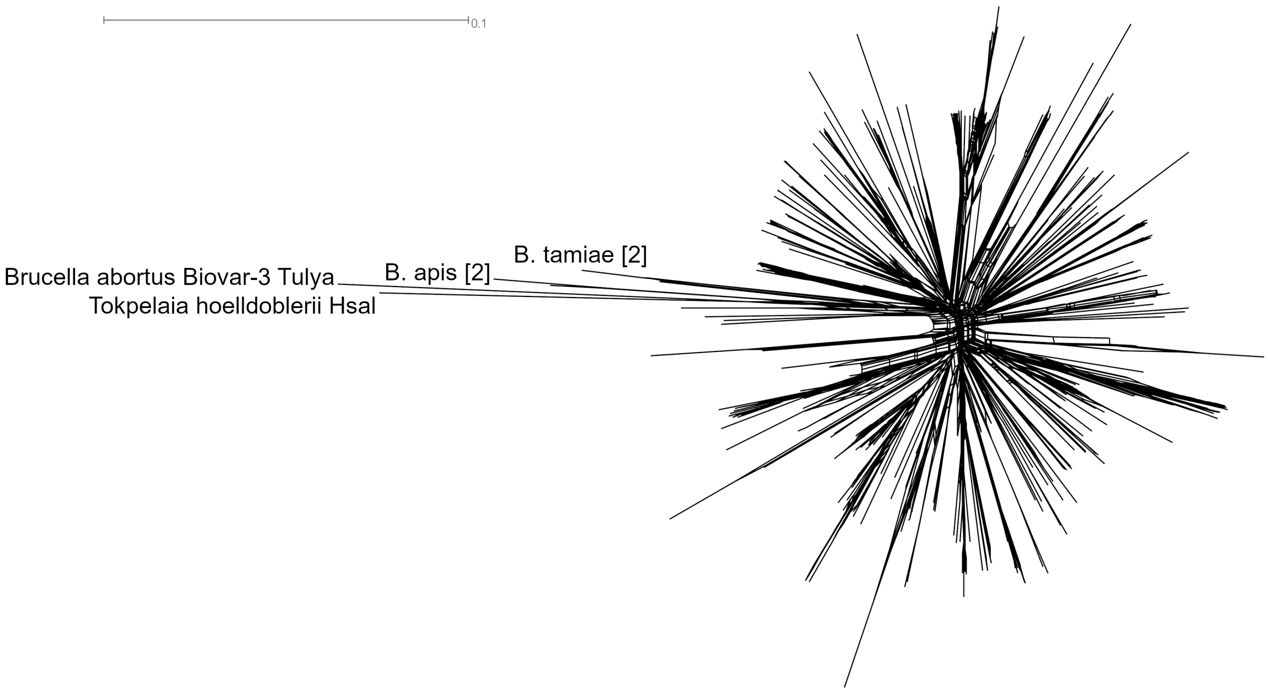


**Fig. S2.** Network phylogeny of *Bartonella* strains. The network was produced using the Neighbor-Net method on uncorrected pairwise distances calculated from an 8345 base pair alignment of nine genetic loci. Distances are show as the number of nucleotide substitutions per site.

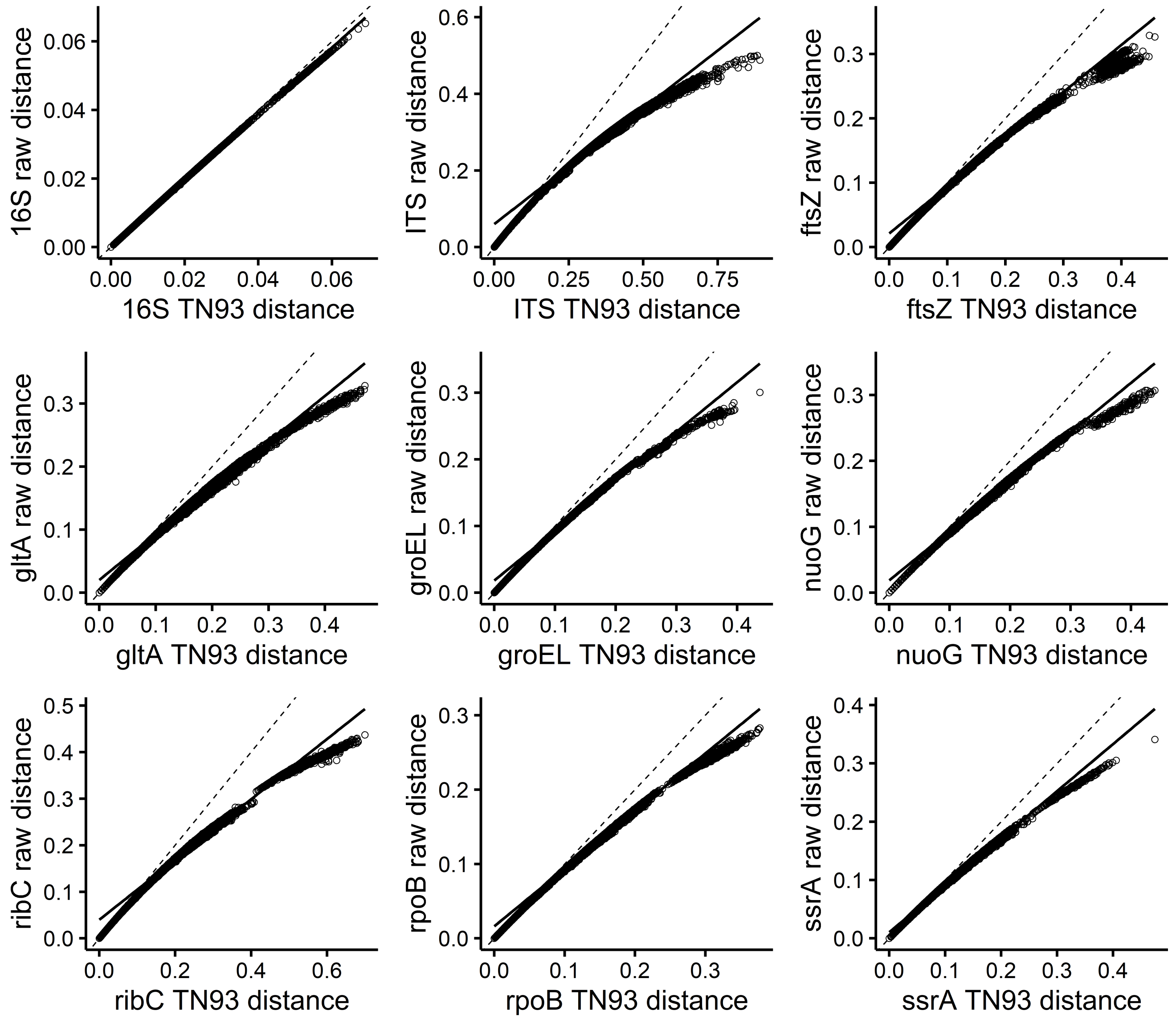


**Fig. S3.** Nucleotide substitution saturation across nine sequenced loci using uncorrected versus adjusted distances. Points represent pairwise distances for all taxa sequenced at each locus. Raw distances represent the uncorrected pairwise distances and adjusted distances were calculated using the Tamura-Nei model. The dashed line shows the 1:1 line for uncorrected versus adjusted distances and the solid line shows the best-fit line for linear regression.

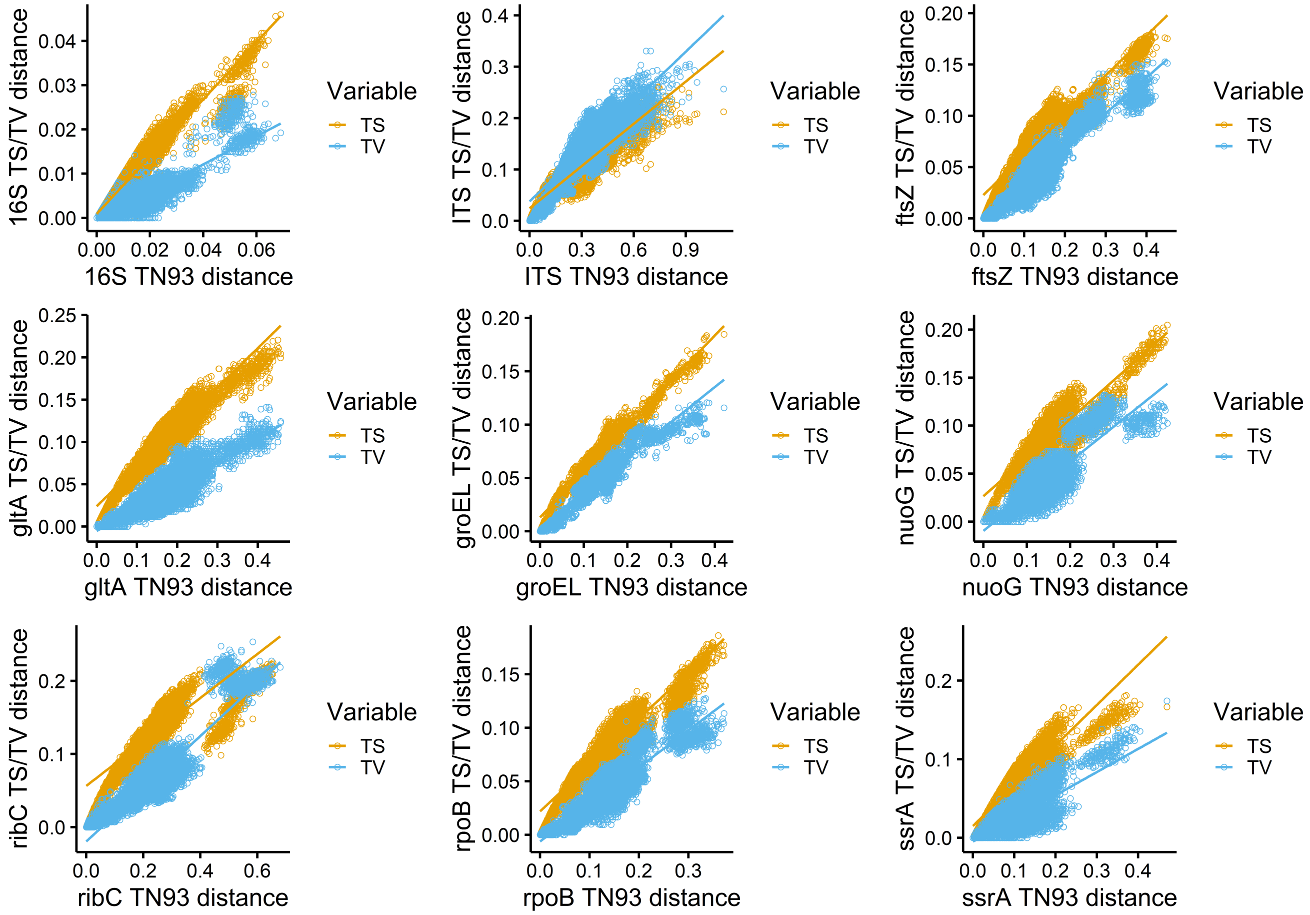


**Fig. S4.** Nucleotide substitution saturation across nine sequenced loci using adjusted distances versus transitions and transversions. Points represent pairwise distances for all taxa sequenced at each locus. Adjusted distances were calculated using the Tamura-Nei model. Transitions (TS) are colored orange and transversions (TV) are colored blue. The solid lines show the best-fit lines for linear regression for transitions and transversions.

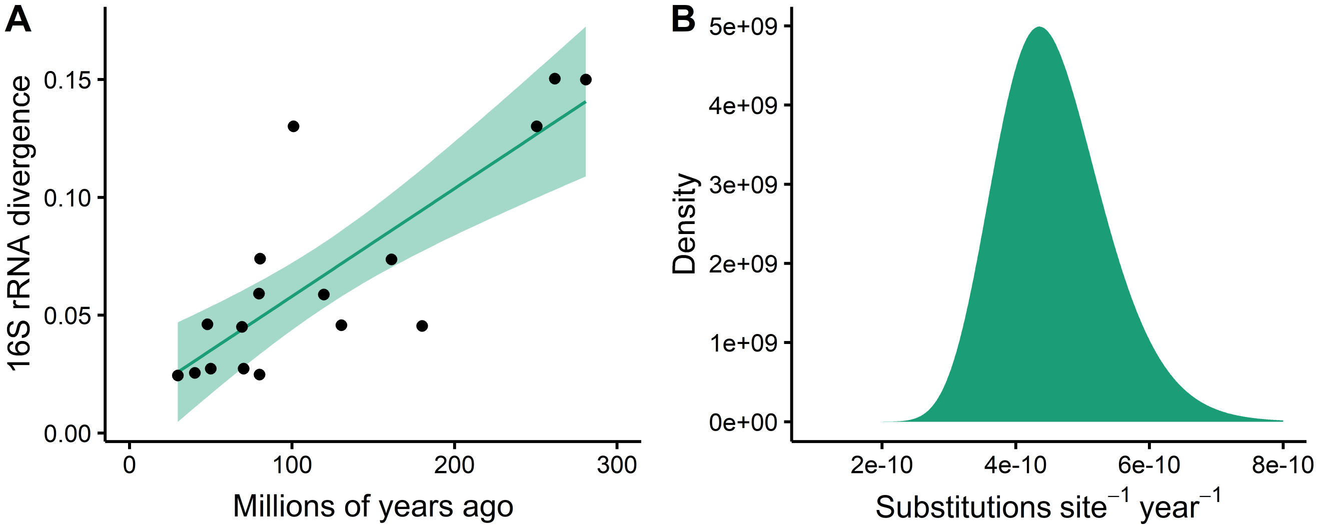


**Fig. S5.** Estimated molecular clock for 16S ribosomal RNA (rRNA). (A) Linear regression of 16S rRNA divergence and host divergence times for bacterial symbionts of arthropods from (70). (B) A lognormal distribution for the 16S rRNA molecular clock estimated by moment matching to the normal distribution of the fitted mean and standard error of the regression.

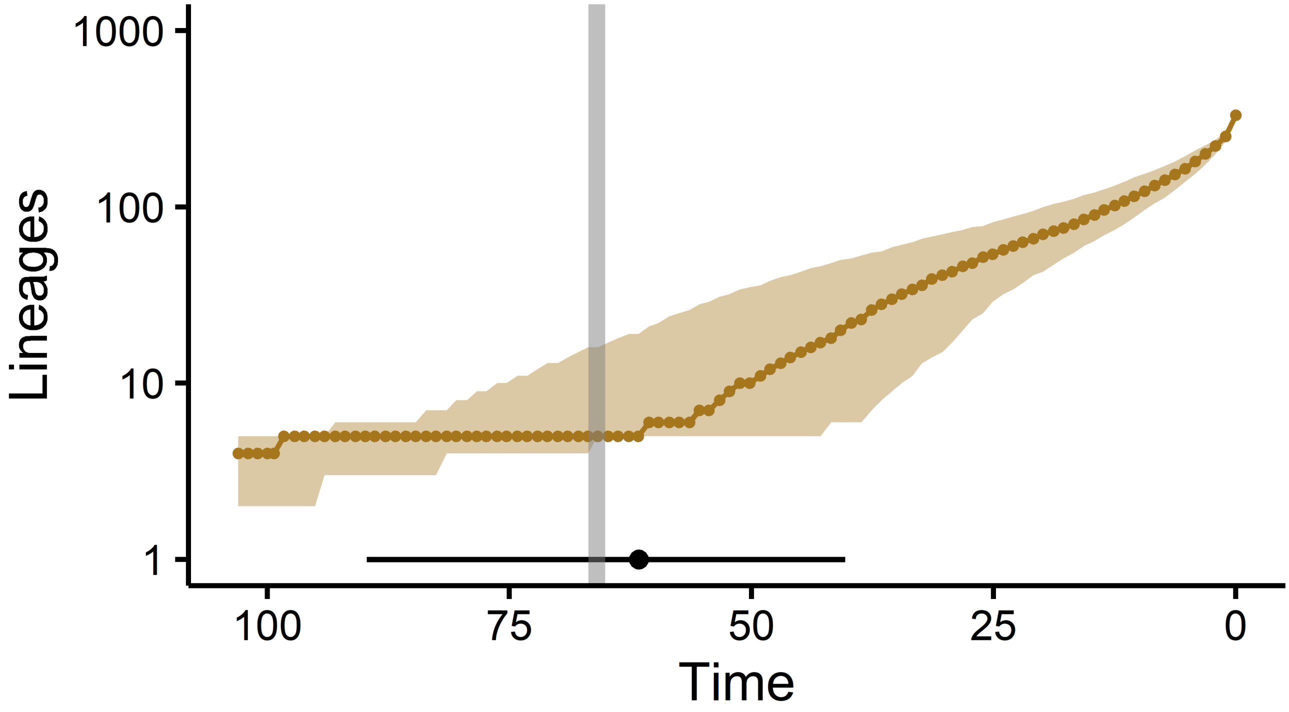


**Fig. S6.** Number of *Bartonella* lineages through time. Circles show the median number of lineages and shading representing the 95% HPD interval. The diversification date of eubartonellae is shown as a circle at the bottom of the figure with a line for the 95% HPD interval. Time is shown in millions of years. The Cretaceous-Paleogene extinction event is drawn as a gray line at 66 million years ago.

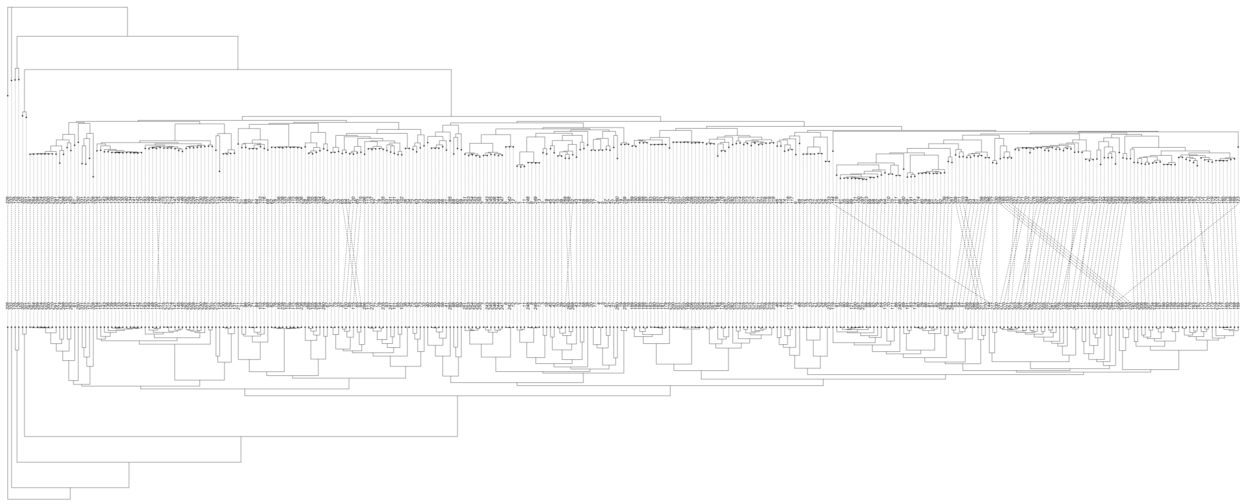

**Fig. S7.** Comparison of Bayesian and maximum likelihood trees. The Bayesian tree (top) used separate sequence evolution models for each of the nine partitioned loci. The maximum likelihood tree (bottom) used concatenated sequences of all nine loci.

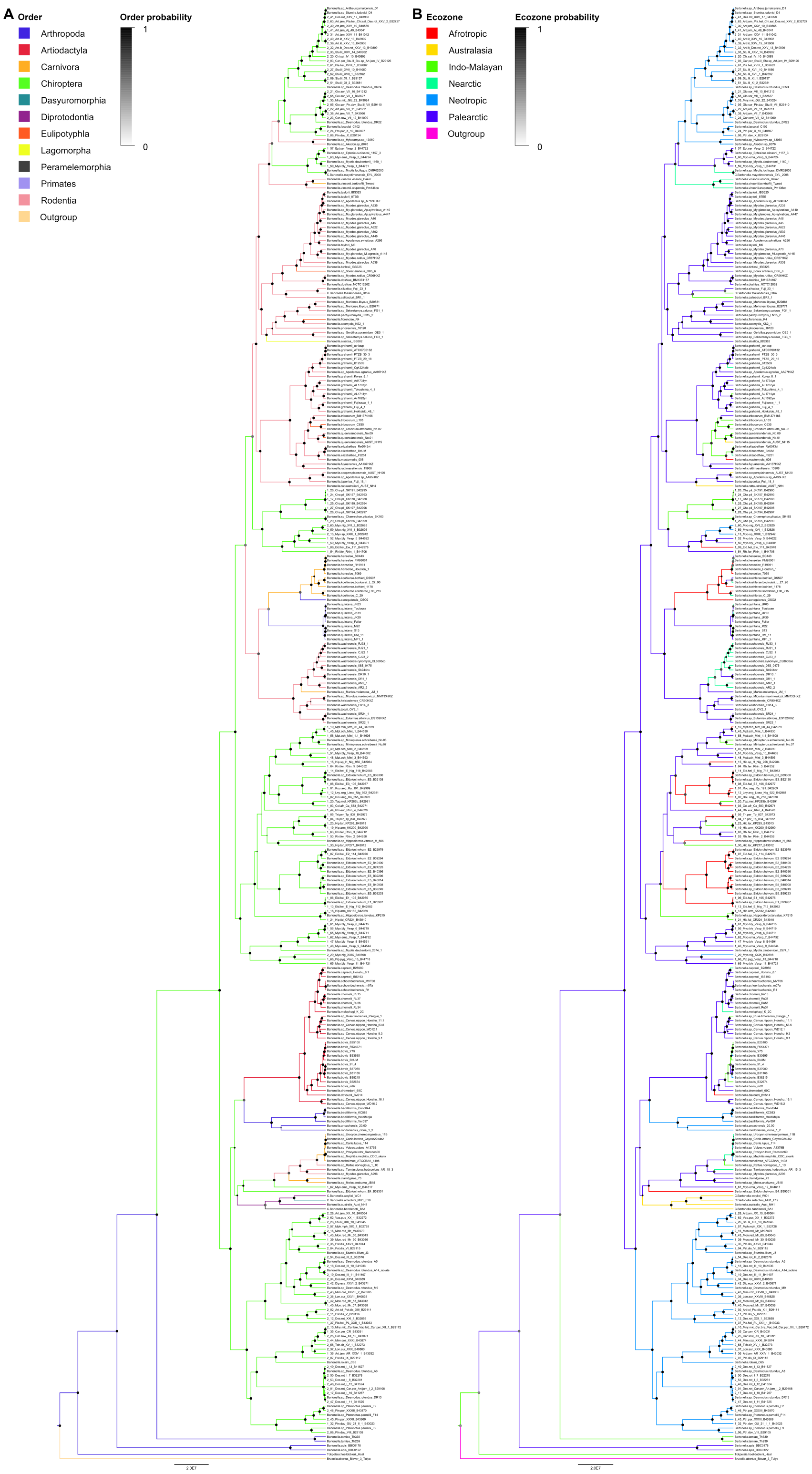


**Fig. S8.** Timed maximum clade credibility tree of *Bartonella* lineages including ancestral reconstruction of (A) host orders and (B) ecozones. Posterior probabilities (PP) for nodes are indicated by the size of circles. Branch lengths are in millions of years. Branches are colored according to their most probable (PP > 0.5) host order or ecozone, with host or ecozone probability shown by the color of circles at each node.

**Table S1.** Features of sequenced genetic loci. The number of taxa with sequences for each locus and the coverage of taxa out of 332 are listed, as well as the number of sites (base pairs) included in the final database. The best DNA substitution model was chosen based on the Akaike information criterion (AIC) in jModelTest. The proportion of invariant sites and the substitution rate gamma shape parameter were estimated from the best model. GTR, generalized time-reversible; TN, Tamura-Nei; G, gamma distributed rate variation; I, proportion of invariant sites.

| Locus | Name | Taxa | Coverage | Sites | AIC best model | Invariant sites | Gamma |
|---|---|---|---|---|---|---|---|
| 16S | 16S ribosomal RNA | 289 | 0.87 | 1511 | GTR+I+G | 0.77 | 0.33 |
| ITS | 16S-23S internal transcribed spacer | 251 | 0.76 | 1833 | GTR+I+G | 0.11 | 1.1 |
| *ftsZ* | cell division protein | 327 | 0.98 | 885 | GTR+I+G | 0.46 | 0.64 |
| *gltA* | citrate synthase | 332 | 1 | 348 | GTR+I+G | 0.32 | 0.71 |
| *groEL* | heat-shock chaperonin protein | 116 | 0.35 | 1632 | GTR+I+G | 0.47 | 0.81 |
| *nuoG* | NADH dehydrogenase gamma subunit | 227 | 0.68 | 342 | GTR+I+G | 0.48 | 0.74 |
| *ribC* | riboflavin synthase | 256 | 0.77 | 561 | GTR+I+G | 0.2 | 0.86 |
| *rpoB* | DNA-directed RNA polymerase beta subunit | 322 | 0.97 | 849 | GTR+I+G | 0.48 | 0.68 |
| *ssrA* | transfer-messenger RNA | 220 | 0.66 | 384 | TN+I+G | 0.31 | 0.6 |

**Table S2.** Results of pairwise homoplasy index (PHI) tests for homologous recombination. The parameter k represents the number of informative sites within a window of 100 base pairs. P-values greater than 0.05 indicate that the observed PHI was outside of the expected distribution of PHI for the tree, thereby failing to reject the null hypothesis of no recombination.

| Locus | k | Expected mean PHI | Expected variance PHI | Observed PHI | P-value |
|---|---|---|---|---|---|
| 16S | 8 | 0.42 | $2.6x10^{-4}$ | 0.423 | 0.57 |
| ITS | 50 | 0.61 | $3.6x10^{-5}$ | 0.78 | 1 |
| *ftsZ* | 43 | 1.81 | $2.5x10^{-4}$ | 1.9 | 1 |
| *gltA* | 57 | 1.24 | $4.9x10^{-4}$ | 1.21 | 0.067 |
| *groEL* | 39 | 1.1 | $3.9x10^{-5}$ | 1.13 | 1 |
| *nuoG* | 45 | 1.62 | $9.5x10^{-4}$ | 1.64 | 0.71 |
| *ribC* | 69 | 1.64 | $2.5x10^{-4}$ | 1.7 | 1 |
| *rpoB* | 43 | 1.93 | $2.9x10^{-4}$ | 1.97 | 0.98 |
| *ssrA* | 45 | 0.63 | $2.2x10^{-4}$ | 0.7 | 1 |
| Concatenated | 40 | 0.95 | $6.5x10^{-6}$ | 1.2 | 1 |

**Table S3.** Results of tests for substitution saturation. The index of substitution saturation (Iss) was calculated for each locus across a series of subtrees randomly pruned to a number of taxa based on 100 iterations. The critical index (Iss.c) is the value at which the sequences will begin to fail to recover the true tree and was calculated for each locus across the series of sampled taxa. If Iss is smaller than Iss.c and the P-value is less than 0.05, then we conclude that the sequences have not experienced severe substitution saturation and can be used for phylogenetic reconstruction. The Iss and Iss.c values shown here assume a symmetrical tree topology and use the proportion of invariant sites from Table S1.

| Locus | Taxa | Iss | Iss.c | T | DF | P-value |
|---|---|---|---|---|---|---|
| 16S | 4 | 0.15 | 0.8 | 8.1 | 24 | 0 |
| | 8 | 0.15 | 0.78 | 7.1 | | 0 |
| | 16 | 0.16 | 0.59 | 4.7 | | 0.0001 |
| | 32 | 0.16 | 0.78 | 6.5 | | 0 |
| ITS | 4 | 0.014 | 1.3 | 76.8 | 21 | 0 |
| | 8 | 0.014 | 1.6 | 82.8 | | 0 |
| | 16 | 0.017 | 0.61 | 26.9 | | 0 |
| | 32 | 0.019 | 2.1 | 85.3 | | 0 |
| *ftsZ* | 4 | 0.33 | 0.82 | 6.1 | 47 | 0 |
| | 8 | 0.33 | 0.82 | 5.6 | | 0 |
| | 16 | 0.33 | 0.58 | 2.7 | | 0.0086 |
| | 32 | 0.34 | 0.85 | 5.6 | | 0 |
| *gltA* | 4 | 0.27 | 0.78 | 12.2 | 121 | 0 |
| | 8 | 0.25 | 0.74 | 11.4 | | 0 |
| | 16 | 0.26 | 0.63 | 8.6 | | 0 |
| | 32 | 0.27 | 0.7 | 10.2 | | 0 |
| *groEL* | 4 | 0.27 | 0.8 | 17.4 | 265 | 0 |
| | 8 | 0.28 | 0.75 | 15.8 | | 0 |
| | 16 | 0.28 | 0.72 | 14.5 | | 0 |
| | 32 | 0.28 | 0.7 | 14 | | 0 |
| *nuoG* | 4 | 0.3 | 0.78 | 10.7 | 117 | 0 |
| | 8 | 0.29 | 0.73 | 9.6 | | 0 |
| | 16 | 0.29 | 0.65 | 7.9 | | 0 |
| | 32 | 0.3 | 0.69 | 8.3 | | 0 |
| *ribC* | 4 | 0.27 | 0.79 | 11.3 | 106 | 0 |
| | 8 | 0.26 | 0.76 | 10.3 | | 0 |
| | 16 | 0.26 | 0.61 | 7.2 | | 0 |
| | 32 | 0.27 | 0.74 | 9.6 | | 0 |
| *rpoB* | 4 | 0.31 | 0.78 | 12.1 | 163 | 0 |

| Locus | Taxa | Iss | Iss.c | T | DF | P-value |
|---|---|---|---|---|---|---|
| | 8 | 0.3 | 0.74 | 10.8 | | 0 |
| | 16 | 0.31 | 0.68 | 9.2 | | 0 |
| | 32 | 0.32 | 0.68 | 9.2 | | 0 |
| *ssrA* | 4 | 0.11 | 1.4 | 15.3 | 13 | 0 |
| | 8 | 0.11 | 1.9 | 18.5 | | 0 |
| | 16 | 0.11 | 0.66 | 5.9 | | 0.0001 |
| | 32 | 0.11 | 2.6 | 26.2 | | 0 |

**Table S4.** Prior distributions for phylogenetic analysis in BEAST.

| Parameter | Distribution | Initial value |
|---|---|---|
| A-C substitutions | gamma(0.05, 10) | 1 |
| A-G substitutions | gamma(0.05, 20) | 1 |
| A-T substitutions | gamma(0.05, 10) | 1 |
| C-G substitutions | gamma(0.05, 10) | 1 |
| G-T substitutions | gamma(0.05, 10) | 1 |
| Base frequencies | uniform(0, 1) | 0.25 |
| Gamma shape parameter | exponential(0.5) | 0.5 |
| Proportion of invariant sites | uniform(0, 1) | 0.5 |
| Birth-death birth rate | uniform(0, 1E5) | 0.01 |
| Birth-death relative death rate | uniform(0, 1) | 0.5 |
| Proportion of taxa sampled | beta(1, 1) | 0.01 |
| 16S rRNA UCED clock rate | lognormal(-21.5, 0.18) | $4.6 \times 10^{-10}$ |
| Host state transition rates | gamma(1, 1) | 1 |
| Ecozone state transition rates | gamma(1, 1) | 1 |

**Table S5.** Robustness of mammal-infecting eubartonellae divergence date to model choice. RelTime divergence dates were estimated in MEGA using uniform prior distributions based on the confidence intervals of 15 host divergence dates listed in Table S7 and a maximum likelihood tree based on a concatenated alignment of all nine loci. BEAST divergence dates were estimated using separate prior distributions for all nine genetic loci separately with a strong prior distribution on the 16S rRNA locus and diffuse continuous-time Markov chain priors on the remaining loci. Separate BEAST runs using alternative sequence evolution, tree, and clock models were run until parameters converged. Intervals in parentheses show either the 95% highest posterior density interval for BEAST analyses or the 95% maximum likelihood confidence interval for RelTime. The primary model used in the main text is in bold. All runs were performed with all nine loci except those marked with a dagger, which were run with ITS. Codon partitioning was added to the last two runs in the table (marked with a double dagger). GTR, generalized time-reversible; TN, Tamura-Nei; I, proportion of invariant sites; G, gamma distributed rate variation; BD, birth-death; BDI, birth-death with incomplete sampling.

| Method | Sequence evolution model | Tree model | Clock model | Divergence date |
|---|---|---|---|---|
| RelTime | GTR+G | Concatenated maximum likelihood | Relative rates | 66.3 (63.5-69.1) |
| BEAST | All loci GTR+I+G | Coalescent, constant size | Strict lognormal | 57.6 (38.1-82.5) |
| BEAST | All loci GTR+I+G | BD | Strict lognormal | 57.2 (37.2-81.7) |
| BEAST | All loci GTR+I+G | BDI | Strict lognormal | 56.9 (35.6-80) |
| BEAST | All loci GTR+I+G | Coalescent, constant size | Relaxed lognormal | 69.6 (45.1-103.1) |
| BEAST | All loci GTR+I+G | BD | Relaxed lognormal | 65.4 (42-96.7) |
| BEAST | All loci GTR+I+G | BDI | Relaxed lognormal | 63.5 (42.3-94.2) |
| BEAST | ssrA TN+I+G, other loci GTR+I+G | BDI | Relaxed lognormal | 63.4 (40.9-97.1) |
| BEAST | All loci GTR+I+G | BDI | Relaxed exponential | **61.6 (40.3-89.7)** |
| BEAST | ssrA TN+I+G, other loci GTR+I+G | BDI | Relaxed exponential | 64.5 (40.8-101.5) |
| BEAST | All loci GTR+I+G | BDI | Relaxed lognormal | 63.4 (41.8-95.8)[†] |
| BEAST | ssrA TN+I+G, other loci GTR+I+G | BDI | Relaxed lognormal | 64 (40.6-101.3)[†] |
| BEAST | All loci GTR+I+G | BDI | Relaxed exponential | 59.9 (39.5-90.3)[†] |
| BEAST | ssrA TN+I+G, other loci GTR+I+G | BDI | Relaxed exponential | 59.5 (37.2-84.1)[†] |
| BEAST | All loci GTR+I+G | BDI | Relaxed lognormal | 67.2 (41.3-97.4)[‡] |
| BEAST | All loci GTR+I+G | BDI | Relaxed lognormal | 65.3 (41.5-97.1)[‡] |

**Table S6.** Summary of *Bartonella* clades and host associations. Host clades above or below the order level associated with each *Bartonella* clade and any named *Bartonella* species or *Candidatus*-level species are listed. Clades A, D, G, L, and N are novel bat-associated clades described in this study. Clade O contains the predominantly rodent-associated clades H-N. Host clades are detailed in Table S6.

| *Bartonella* clade | Tips in clade | Host order(s) | Host clade | *Bartonella* species in clade |
|---|---|---|---|---|
| A | 51 | Chiroptera | Noctilionoidea | *Candidatus* B. rolaini |
| B | 4 | Dasyuromorphia<br>Diprotodontia<br>Peramelemorphia | Marsupialia | *B. australis* |
| | | | | *Candidatus* B. antechini |
| | | | | *Candidatus* B. bandicootii |
| | | | | *Candidatus* B. woyliei |
| C | 32 | Artiodactyla | Pecora | *B. bovis* |
| | | | | *B. capreoli* |
| | | | | *B. chomelii* |
| | | | | *B. dromedarii* |
| | | | | *B. melophagi* |
| | | | | *B. schoenbuchensis* |
| | | | | *Candidatus* B. davousti |
| D | 55 | Chiroptera | Yinpterochiroptera | *B. naantaliensis* |
| E | 19 | Rodentia | Sciuridae | *B. jaculi* |
| | | | | *B. heixiaziensis* |
| | | | | *B. washoensis* |
| F | 10 | Carnivora | Felidae | *B. henselae* |
| | | | | *B. koehlerae* |
| G | 15 | Chiroptera | Vespertilionoidea | |
| H | 29 | Rodentia | Murinae | *B. elizabethae* |
| | | | | *B. fuyuanensis* |
| | | | | *B. grahamii* |
| | | | | *B. queenslandensis* |
| | | | | *B. rattimassiliensis* |
| | | | | *B. mastomydis* |
| | | | | *B. tribocorum* |
| I | 4 | Rodentia | Gerbillinae | *B. pachyuromydis* |
| J | 22 | Rodentia | Arvicolinae | *B. birtlesii* |
| | | | | *B. doshiae* |
| | | | | *B. taylorii* |
| K | 3 | Rodentia | Neotominae | *B. vinsonii* |

| *Bartonella* clade | Tips in clade | Host order(s) | Host clade | *Bartonella* species in clade |
|---|---|---|---|---|
| L | 7 | Chiroptera | Myotis | *Candidatus* B. mayotimonensis |
| M | 2 | Rodentia | Sigmodontinae | |
| N | 30 | Chiroptera | Phyllostomidae | |
| O | 88 | Rodentia | Muroidea | |

**Table S7.** Description of *Bartonella* clades and host associations. MRCA, most recent common ancestor.

| *Bartonella* clade | Host clade | Description |
|---|---|---|
| A | Noctilionoidea | MRCA for families Noctilionidae, Mormoopidae, and Phyllostomidae in order Chiroptera |
| B | Marsupialia | MRCA for orders Dasyuromorphia, Diprotodontia, and Peramelemorphia in infraclass Marsupialia |
| C | Pecora | MRCA for families Bovidae and Cervidae in order Artiodactyla |
| D | Yinpterochiroptera | MRCA for families Rhinolophidae, Hipposideridae, and Pteropodidae in order Chiroptera |
| E | Sciuridae | MRCA for genera *Sciurus*, *Tamiasciurus*, *Glaucomys*, *Eutamias*, *Urocitellus*, *Spermophilus*, *Otospermophilus*, and *Cynomys* in family Sciuridae |
| F | Felidae | MRCA for genera *Panthera*, *Lynx*, *Felis*, *Puma*, and *Acinonyx* in family Felidae |
| G | Vespertilionoidea | MRCA for families Vespertilionidae and Molossidae in order Chiroptera |
| H | Murinae | MRCA for genera *Bandicota*, *Rattus*, *Niviventer*, *Melomys*, *Uromys*, and *Mus* in subfamily Murinae |
| I | Gerbillinae | MRCA for genera *Pachyuromys*, *Meriones*, and *Sekeetamys* in subfamily Gerbillinae |
| J | Arvicolinae | MRCA for subfamily Arvicolinae |
| K | Neotominae | MRCA for subfamily Neotominae |
| L | *Myotis* | MRCA for species *Myotis blythii* and *Myotis lucifugus* in family Vespertilionidae |
| M | Sigmodontinae | MRCA for genera *Hylaeamys*, |

| | | |
|---|---|---|
| | | *Akodon*, and *Sigmodon* in subfamily Sigmodontinae |
| N | Phyllostomidae | MRCA for family Phyllostomidae |
| O | Muroidea | MRCA for superfamily Muroidea (excluding *genus* Typhlomys) |

**Table S8.** Comparison of divergence date estimates for 15 *Bartonella* clades with divergence dates of the associated hosts within each clade collated from TimeTree. The number of published studies used to estimate host divergence dates is listed. Both host and *Bartonella* clade divergence dates are in units of millions of years. Intervals in parentheses show either the 95% highest posterior density interval for *Bartonella* clade dates or the 95% confidence interval for host clade dates. Intervals in brackets show the ranges. Details regarding *Bartonella* clades are found in Tables S5-S6. The positive correlation between *Bartonella* and host clades is depicted in Fig. 2.

| *Bartonella* clade | TimeTree studies | Host clade date | *Bartonella* clade date |
|---|---|---|---|
| A | 19 | 43 (41-46) [36.7-60.4] | 45.9 (29.6-68.4) [24.1-110.1] |
| B | 17 | 62 (58-67) [49.8-82] | 35.1 (21.9-53.8) [16-84.7] |
| C | 10 | 27.3 (23.1-31.5) [20.8-38.7] | 15.4 (9.4-23.1) [6.9-36.2] |
| D | 21 | 58 (56-61) [46-71.2] | 49.6 (32.3-72.5) [25.2-110.1] |
| E | 11 | 35 (29-40) [17.8-47.6] | 18.8 (10.7-28.9) [8.7-52.9] |
| F | 12 | 15.2 (12.3-18.1) [9.6-26.3] | 8.4 (4.8-13) [3.3-27.8] |
| G | 15 | 49 (45-52) [36-60.4] | 36.9 (22.9-55.1) [18.7-84.3] |
| H | 84 | 20.9 (18.3-23.4) [8.8-53.6] | 20.8 (13.1-30.6) [9.7-47.5] |
| I | 6 | 18.4 (10.3-26.4) [11-28.4] | 16.4 (9.1-25.1) [6.9-38.5] |
| J | 3 | 18.6 [15.2-20.9] | 25.2 (15.5-37.1) [11.7-47.2] |
| K | 8 | 19.3 (12.1-26.4) [8.6-32] | 11.3 (6.1-18.6) [4.2-35.4] |
| L | 6 | 18.1 (9.3-27) [10.8-32.8] | 10.5 (5.7-17.2) [4.7-31.9] |
| M | 5 | 19.8 (10-29.5) [11.6-29.7] | 7.4 (2.7-15.2) [1.6-30.7] |
| N | 16 | 31 (29-33) [25-35.3] | 23.9 (15.5-35.2) [11.9-54.4] |
| O | 16 | 45 (42-49) [35.9-60.1] | 40.4 (26.4-59) [21.1-95] |

**Table S9.** Posterior median estimates of clock rates across genetic loci. Numbers in parentheses show the 95% highest posterior density (HPD) interval. UCED, uncorrelated exponential distribution. Median UCED clock rate represents the molecular clock rate (x$10^{-9}$ substitutions site$^{-1}$ year$^{-1}$). Median branch clock rate represents the molecular clock rate estimate weighted by branch lengths (x$10^{-9}$ substitutions site$^{-1}$ year$^{-1}$).

| Locus | Median UCED clock rate | Median branch clock rate |
|---|---|---|
| 16S | 0.47 (0.31-0.63) | 0.52 (0.34-0.71) |
| ITS | 9.2 (5.9-13.3) | 11.5 (7.5-16.6) |
| *ftsZ* | 3.1 (2-4.6) | 3.8 (2.4-5.4) |
| *gltA* | 3.8 (2.3-5.5) | 3.7 (2.4-5.3) |
| *groEL* | 2.5 (1.6-3.7) | 2.4 (1.5-3.5) |
| *nuoG* | 3.3 (2-4.8) | 3.3 (2.1-4.8) |
| *ribC* | 4 (2.4-5.6) | 3.9 (2.5-5.6) |
| *rpoB* | 5 (3.1-7.1) | 4.8 (3-6.7) |
| *ssrA* | 2.9 (1.8-4.2) | 2.8 (1.7-4.1) |

**Table S10.** Tip-association tests of host trait clustering on trees. Observed credible intervals were drawn from 1000 posterior sampled trees. Null distributions were produced from 100 resampling steps for each sampled tree. ML, maximum likelihood; AI, association index; PS, parsimony score.

| | Posterior sampled trees | | Single ML tree | |
|---|---|---|---|---|
| Trait | Order | Ecozone | Order | Ecozone |
| States | 12 | 7 | 12 | 7 |
| Observed AI | 1.4 (1.39-1.43) | 6.13 (5.87-6.25) | 2.2 | 6.1 |
| Null AI | 25.5 (23.1-27.4) | 28.1 (25.8-30) | 20.9 (19-22.8) | 23 (21.2-24.9) |
| Observed PS | 24 (24-24) | 61.9 (61-62) | 54 | 102 |
| Null PS | 153.5 (147-160) | 178.4 (172.5-186.1) | 172.2 (148-205) | 193.8 (175-216) |

**Table S11.** Posterior probability of host and ecozone states for the mammal-infecting eubartonellae ancestor. For the MCC tree, 100 stochastic character mapping simulations were run on 100 randomly resampled trees and for the posterior sampled trees, 100 stochastic simulations were run on 10 randomly chosen trees with 10 random resampling iterations of tips. The distribution of the posterior probability of the ancestral state over 100 trees is summarized by the median and the interquartile range (in parentheses).

| | Sampled tips | Probability host is Chiroptera | Probability ecozone is Palearctic |
|---|---|---|---|
| MCC tree | 87 | 0.99 (0.95-1) | 0.75 (0.7-0.8) |
| | 32 | 0.95 (0.91-0.98) | 0.63 (0.56-0.69) |
| | 21 | 0.93 (0.88-0.96) | 0.64 (0.56-0.69) |
| Posterior sampled trees | 87 | 0.99 (0.95-1) | 0.77 (0.7-0.82) |
| | 32 | 0.92 (0.87-0.95) | 0.67 (0.55-0.74) |
| | 21 | 0.93 (0.9-0.97) | 0.63 (0.57-0.72) |

**Table S12.** Results of stochastic character mapping of host orders and ecozones on 1000 posterior sampled trees. The posterior distribution of the number of transitions is given as the median and the 95% HPD interval (in parentheses).

| Network | Transition | Count |
|---|---|---|
| Order | Arthropoda → Chiroptera | 1 (0-1) |
| | Arthropoda → Outgroup | 1 (0-1) |
| | Carnivora → Arthropoda | 1 (0-1) |
| | Chiroptera → Arthropoda | 1 (0-2) |
| | Chiroptera → Artiodactyla | 1 (0-1) |
| | Chiroptera → Carnivora | 1 (0-2) |
| | Chiroptera → Diprotodontia | 1 (0-2) |
| | Chiroptera → Rodentia | 2 (1-4) |
| | Diprotodontia → Dasyuromorphia | 1 (0-1) |
| | Diprotodontia → Peramelemorphia | 1 (0-1) |
| | Rodentia → Carnivora | 3 (2-5) |
| | Rodentia → Chiroptera | 2 (1-3) |
| | Rodentia → Eulipotyphla | 4 (3-4) |
| | Rodentia → Lagomorpha | 1 (0-1) |
| | All order transitions | 26 (24-30) |
| Ecozone | Afrotropic → Indo-Malayan | 2 (0-5) |
| | Afrotropic → Nearctic | 3 (1-5) |
| | Afrotropic → Palearctic | 3 (0-7) |
| | Australasia → Indo-Malayan | 1 (0-2) |
| | Australasia → Palearctic | 1 (0-3) |
| | Indo-Malayan → Afrotropic | 2 (0-4) |
| | Indo-Malayan → Australasia | 1 (0-2) |
| | Indo-Malayan → Nearctic | 1 (0-3) |
| | Indo-Malayan → Neotropic | 2 (0-3) |
| | Indo-Malayan → Palearctic | 3 (1-5) |
| | Nearctic → Afrotropic | 1 (0-4) |
| | Nearctic → Indo-Malayan | 1 (0-3) |
| | Nearctic → Neotropic | 1 (0-3) |
| | Nearctic → Palearctic | 4 (2-7) |
| | Neotropic → Afrotropic | 1 (0-3) |
| | Neotropic → Indo-Malayan | 1 (0-3) |

| Network | Transition | Count |
|---|---|---|
| | Neotropic → Palearctic | 2 (0-5) |
| | Outgroup → Indo-Malayan | 1 (0-2) |
| | Palearctic → Afrotropic | 8 (4-12) |
| | Palearctic → Australasia | 3 (1-6) |
| | Palearctic → Indo-Malayan | 12 (9-16) |
| | Palearctic → Nearctic | 11 (8-14) |
| | Palearctic → Neotropic | 6 (3-9) |
| | Palearctic → Outgroup | 1 (0-3) |
| | All ecozone transitions | 82 (71-92) |

**Table S13.** Node properties of state transition networks. Measures are based on median counts of stochastic character mapping simulations on 1000 posterior sampled trees. The networks exclude transitions between states and the outgroup (*Brucella abortus*) or transitions between mammalian orders and arthropods.

| Network | State | Degree | Weighted degree | Out-degree | Weighted out-degree | Betweenness |
|---|---|---|---|---|---|---|
| Order | Artiodactyla | 1 | 1 | 0 | 0 | 0 |
| | Carnivora | 2 | 4 | 0 | 0 | 0 |
| | Chiroptera | 5 | 7 | 4 | 5 | 4 |
| | Dasyuromorphia | 1 | 1 | 0 | 0 | 0 |
| | Diprotodontia | 3 | 3 | 2 | 2 | 4 |
| | Eulipotyphla | 1 | 4 | 0 | 0 | 0 |
| | Lagomorpha | 1 | 1 | 0 | 0 | 0 |
| | Peramelemorphia | 1 | 1 | 0 | 0 | 0 |
| | Rodentia | 5 | 12 | 4 | 10 | 2 |
| Ecozone | Afrotropic | 7 | 20 | 3 | 8 | 0.33 |
| | Australasia | 4 | 6 | 2 | 2 | 0 |
| | Indo-Malayan | 10 | 26 | 5 | 9 | 3.67 |
| | Nearctic | 7 | 22 | 4 | 7 | 0.33 |
| | Neotropic | 6 | 13 | 3 | 4 | 0 |
| | Palearctic | 10 | 53 | 5 | 40 | 3.67 |
